# Multifunctional evolution of palm squirrel coat colour and pattern

**DOI:** 10.1101/2023.09.15.557893

**Authors:** M Nivetha, Senan D’Souza, AC Shijisha, Russell A. Ligon, R Nandini

**Affiliations:** Indian Institute of Science Education and Research Tirupati, India; Cornell University, Ithaca, New York, United States of America

## Abstract

Complex colour pattern traits often evolve in response to multiple selective pressures, emphasizing the need for their investigation in an integrative framework. We use museum specimens (n=718) to study the evolution of coat colouration in three palm squirrel species (*Funambulus* spp.) in the Indian subcontinent, with climate, vegetation, and predators as the potential selective forces. Our results indicate a strong geographical trend in coat colour distribution within and across species, with darker individuals in the southern and lighter individuals in the northern regions. We find significant darkening of coat colouration in more humid, densely vegetated regions with dark soils. Fine-scale intra-specific analyses suggest that species respond differently to multiple abiotic selective pressures. Investigation of stripe pattern elements suggests that the colour-based elements are more visible to raptors, while luminance-based elements are more visible to ferrets. We also find that squirrel species are darker in areas with denser and taller canopies, with low-contrast stripes specific to each predator’s vision. Overall, squirrel coat colouration evolves in response to multiple abiotic factors, including climate and habitat, and specific stripe elements provide particular visual effects on predators depending on the habitat.

## Introduction

Morphological traits like colouration perform multiple adaptive functions in response to ecophysiological stressors, with implications at various biological levels [1,2]. Colour and pattern traits across an animal’s body can be simple or complex depending on the diversity and nature of the trait elements. In addition, different selective pressures may shape an animal’s overall phenotype and its individual elements [3]. An animal’s overall visual appearance could be an optimal or simultaneous response to multiple, mutually non-exclusive stressors acting on its different elements [4]. However, most studies investigating colour and pattern evolution focus on individual elements and functions. This study explores the evolution of coat colour and patterns in a genus of small diurnal squirrels in the Indian subcontinent as concurrent responses to abiotic and biotic selective pressures. Coat colouration and patterns in mammals could have evolved for inter-sexual signaling within a species, environmental adaptation, or inter-specific signaling (e.g., predator and prey)[2].

Colouration is a significant sexual signal in birds with sexual dimorphism, where males are typically more colourful [5]. Few studies have addressed mammalian colouration as a sexual signal, except in a few sexually dichromatic primate species [2,6–8]. Even in polygynous mammalian systems, sexual dichromatism is rare, suggesting the relative insignificance of colouration in sexual signaling compared to birds [9].

The most well-studied explanation for animal colouration is its evolution in response to abiotic selective pressures, including climate and habitat variables [10–12]. Adaptation to the environment could serve camouflage and physiological functions. According to Gloger’s rule, darker animals are present in warm and humid areas [13]. Increased precipitation and vegetation result in darker environments with low light conditions, where darker animals can camouflage better. Populations of *Peromyscus* rodents evolved colouration that helps camouflage themselves in response to specific elements of their abiotic environments, like soil colouration [14]. Adaptations to climatic conditions such as temperature, solar radiation, and humidity also provide physiological advantages, like regulating heat influx and protecting the animal from ultraviolet (UV) radiation [13,15,16]. Lighter colours absorb less heat, and darker colours are better at protecting from UV damage [17]. Hence, in the case of physiological functions such as thermoregulation and UV protection, we expect lighter colouration in warmer temperatures and darker colouration in regions with high solar radiation. The responses to abiotic and biotic selective pressures often could be interactive and not necessarily exclusive.

Camouflage as a coat colour adaptation could be achieved by multiple mechanisms, including crypsis and disruptive colouration [18,19]. Both plain and patterned colouration on an individual’s body could be cryptic depending on the complexity levels of background surfaces. Regardless of an animal’s background surface, distinct contrasting markings on the inside or at the edges of its body can break down its outline, making it less visible to predators [20–22]. In such cases, the characteristics of pattern elements, such as the position of markings on the animal’s body, optimal contrasts, and intensity of the edges between markings or patches, determine the effectiveness of camouflage by disruptive colouration [23,24]. Mechanisms like edge detection, feature disruption, pictorial relief, and differential grouping help patterns achieve visually disruptive effects [25–28]. Patterns such as contrasting longitudinal stripes confer motion dazzle effects, which make being spotted by predators hard while the prey moves [24,29–32].

Although contradictory results are available regarding the efficiency of high-contrast stripes over low-contrast ones [24,33], recent simulation studies with human predators controlling the size, contrast, orientation, and movement properties of targets revealed that the presence of high internal contrast on smaller targets moving at moderate speed had the best efficiency at motion dazzle [85]. Also, parallel stripes underestimate the speed of target movement, while perpendicular stripes overestimate it. Because the great conspicuity of the markings in larger prey negates any benefit from the motion dazzle effect, motion dazzle marks may be more likely to evolve in smaller animals [34].

The visual systems of predators often determine the evolution of colour and pattern traits of prey species [35]. For instance, pattern elements could provide different visual outcomes for aerial and terrestrial predators, given the enormous variation in their visual sensitivities [36].

Experimental studies have shown disruptive colouration functions by exploiting the edge-detection ability of viewers [30,37,38]—edge detection functions by detecting contours and boundaries between contrasting colour or luminance patches. While most visual systems can detect contrasts and edges [39], their differential sensitivities to the properties of elements like chromaticity and luminance that constitute the edges remain unclear.

Because colour and patterns provide adaptive functions across abiotic and biotic ecophysiological selective pressures, studying colour and pattern evolution requires an integrative study framework [40]. Here, we investigate the evolution of palm squirrel colour, pattern evolution in response to abiotic selective pressures such as climate and habitat, and biotic pressures from predators with different visual systems.

Palm squirrels (*Funambulus* spp.) are endemic to the Indian subcontinent, with three out of six species occupying most of the geographical range of the genus, including a wide variety of climatic and habitat conditions [41]. Their habitat and climatic ranges include shrublands and dry areas *(Funambulus pennantii)*, deciduous and scrub forests (*F. palmarum)*, and evergreen rainforests *(F. tristriatus)* [42]. The phylogenetic relationship among these three species is currently incomplete. However, the number of chromosomes and physical and acoustic similarities (ongoing work) suggest that *F. tristriatus* could be more closely related to *F. palmarum* than *F. pennantii*.

The most distinguishing feature of palm squirrels is their distinct longitudinal (parallel to body length) alternating dark and light stripes on their dorsal and lateral (for *F. pennantii* only) bodies, implying that apart from having a complex body pattern for background matching, longitudinal stripes could perform anti-predatory functions through disruptive and motion dazzle effects. The embryonic development, genetic mechanisms, and evolution of rodent stripes have been studied previously [43]. However, while many rodents and other small mammals display longitudinal stripes on their dorsal and lateral bodies, their adaptive significance in anti-predator mechanisms accounting for predator visual sensitivities needs investigation. Given the extensive geographic distribution of palm squirrels, there could be local adaptations depending on specific levels of exposure to selective pressures. Moreover, palm squirrels occupy broad habitat classes, with predation pressures from aerial and terrestrial predators across bird and mammalian taxa, making them an excellent system to investigate coat colour adaptation determined by multiple biotic and abiotic selective pressures.

The questions we address in this study are:

1. How does squirrel colour vary across geography?
2. Did palm squirrel coat colouration evolve as a response to climate and habitat?
3. Did squirrels evolve stripe patterns in response to predators with different visual capacities?
4. Does squirrel colour and pattern covary with forest canopy properties?

While a few studies have examined the evolution of coat colour in squirrels [44,45], this is the first study to investigate the evolutionary forces driving the tree squirrel coat colour and pattern distribution in India.

## Methods

### Sample details and museum skins photography

We conducted the study by examining museum skins of *Funambulus* squirrels at the Bombay Natural History Society (BNHS) Museum, Mumbai, India. The specimens were collected from 1906 to 1969 from across the Indian subcontinent, with representatives from most of each species’ geographical distribution (Figure 1). We obtained specimen details such as body morphometrics, age, sex, and collection date from the specimen tags.

**Figure 1.**
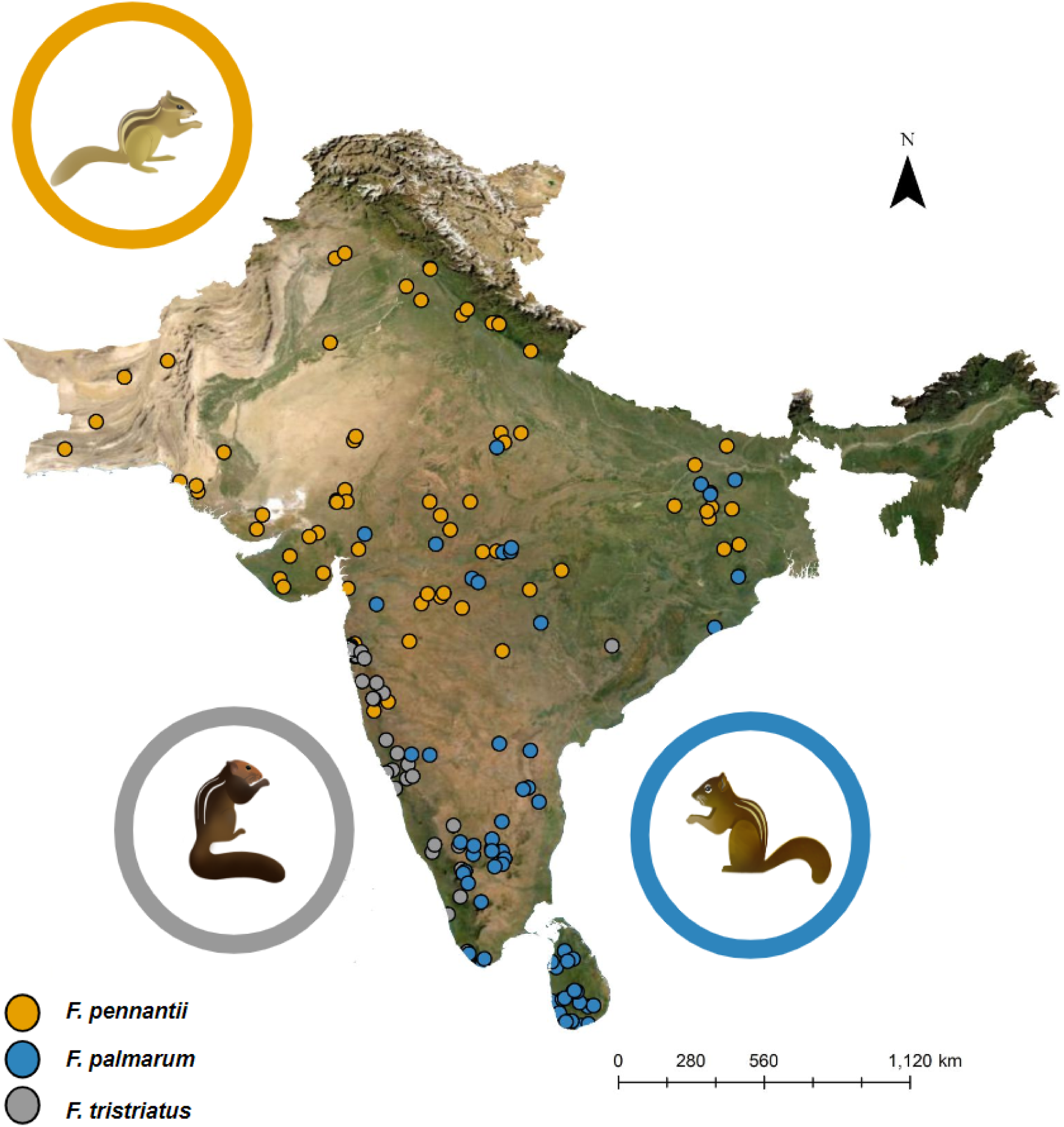
Study area map. Description: Map showing the locations of squirrel specimen collection, including *F. palmarum* (Indian palm squirrel; blue circles; n=204), *F. pennantii* (Northern palm squirrel; yellow circles; n=397), and *F. tristriatus* (Jungle palm squirrel; grey circles; n=114).

We used a light-controlled photo studio box and a Nikon D3400 DSLR camera with an 18-55mm kit lens for photographing the specimens. We maintained constant exposure and aperture settings, and in addition, we placed a ruler and a gray card in each frame for size and exposure control (Supplementary Figure S1). We photographed 718 specimens, including all three species (386 *F. pennantii*, 216 *F. palmarum,* and 116 *F. tristriatus*), in RAW format from three angles (dorsal, left, and right lateral). Only the right lateral images of the squirrel were used for the lateral image analysis to avoid complications of asymmetry, if any.

### Colour quantification from select body regions

We generated multispectral images from the RAW photographs of squirrels using the micaToolbox [46] plugin in ImageJ [47] with an 18% gray card as the standard. We then manually drew polygons over the major body regions, including dark stripes, light stripes, body(outside the stripes–where the forearm starts), head, and neck (Supplementary Figure S2) across a total of ten regions (four dark stripes, three light stripes, one base body, one neck, and one head). The reflectance of a particular body region in the R, G, and B channels was estimated from individual selection polygons in each specimen (Supplementary Figure S3) using the batch multispectral image analysis tool in micaToolbox with average scale values and the FTT framework [48].

### Cone-catch models to simulate predator vision

We quantified colour and stripe parameters under small mammalian carnivore and raptor visual systems, as these are the primary predators of squirrels. We used the in-built ferret cone-catch model in micaToolbox. For the raptor visual model, we created a cone-catch model using extracted sensitivity data for spectra including violet, longwave, mediumwave, shortwave, and double cones (luminance channel) [40, 41] using the WebPlot digitizer [49]. We customized both predator cone-catch models to our camera model using the X-Rite colour Checker Passport [50] under D65 illumination following the micaToolbox pipeline.

### Dorsal and lateral longitudinal stripe pattern quantification

We used the Quantitative Colour and Pattern Analysis (QCPA) framework [46] from micaToolbox to estimate the pattern and complexity parameters for the dorsal and lateral body stripes. Firstly, we generated multi-spectral images from the RAW images by marking a region of interest encompassing the stripes. A selection of approximately 1.5 inches^2^, including the four dark and three light stripes, was created for all the dorsal images. For *F*. *palmarum* and *F*. *tristriatus,* a square of 1 cm^2^ was selected across the lateral images, starting from the lateral dark stripe and ending with the flank region. For *F. pennantii,* which has five stripes, this selection included the lateral light stripe, but we maintained the dimension of the ROI constant across species (Supplementary Figure S4).

After removing negative values, we converted the multispectral images with the selection into ferret and raptor cone-catch models. We used the double-cone channel for raptor vision and the longwave channel for ferret vision for processing luminance, as most dichromats visualize achromatic information through the longwave channel [51]. We ran the QCPA framework with Gaussian acuity correction, RNL ranked filter clustering, and a luminance Weber fraction 0.1 for both raptor and ferret model pipelines. We used the Weber fractions associated with different channels under raptor vision (0.1: 0.05 : 0.05: 0.025), as suggested by [52]. We set the acuity value at 30 degrees per cycle for the raptor cone-catch model and 10 degrees per cycle for the ferret cone-catch model. The viewing distance was made constant at 500mm across both visual models to maximize the number of output parameters; as for ferret vision, increasing the distance resulted in several ‘NA’ values in the output. We rescaled the images to 5 pixels per MRA and proceeded by selecting the region of interest. We set the colour and luminance JND thresholds at three and all other parameters at default. We generated colour pattern parameters under three broad analyses - colour adjacency analysis, Visual contrast analysis [53], Boundary strength analysis [46], and basic cluster statistics.

In addition, we sequentially measured 27 measurements of the length and width of both dark and light stripes from dorsal and lateral images at three locations across the length of the body in ImageJ [47]. Due to the high correlation between variables, we investigated species-level differences in eight length and width measurements relative to body size.

### Extraction of environmental variables from sampling locations

We manually extracted geographical coordinates using the location details from museum collections using Google My Maps. We downloaded 25 environmental variables (Supplementary Table S1) from various online sources. We resampled all variables to a spatial resolution of 10km from their original resolution using ArcMap 10.5 [54]. We used this coarse resolution due to the low accuracy of coordinates of the collection locations, resulting from the availability of only broad location information for many museum records (village/town level). We grouped variables into five major categories - precipitation, temperature, solar radiation, vegetation, and soil characteristics to delineate the effects of multiple abiotic stressors on coat colouration. We calculated relative humidity from dewpoint temperature and surface pressure [55]. We calculated the standard deviation of a year’s temperature and precipitation data to measure a location’s seasonality. Wherever possible, we used the earliest available raster layers (1979-1980) for each variable to represent climate and habitat features prevalent at the time of specimen collection. In case of the non-availability of layers for this period, we used the earliest available data. We checked all variables for correlations and used only the less correlated (<70 percent) variables for further analysis (Supplementary Figure S5).

### Statistical analyses

We performed all downstream statistical analyses in R [56]. We inspected data distribution using the R package fitdistplusr [57]. We performed separate principal component analyses using reflectances from RGB channels for each species and each body region to estimate intraspecies and interspecies colour variation, respectively. We used the cumulative reflectance score of the first principal component (over 90% for pooled and species-subset data) for further analysis. Hereafter, we refer to this metric as colour. After removing body colour outliers, we retained 643 skins for the rest of the analyses. We performed ANOVA and post hoc Tukey-HSD tests for each body region to test a) the colour variation within specific body parts across species and b) the colour variation across body regions within species. All further analyses are conducted using the body colour metric and will be referred to as coat colour from here onwards.

We performed hierarchical generalized additive models using the mgcv package [58] in R to predict coat colour variation across geography. We used the two-way interactive effects of latitude and longitude as smoothing parameters (thin plate regression splines) for the overall data with species as the grouping factor. In addition, we used altitude with species-level grouping factors [59]. Based on the best model, we then predicted the coat colour of individuals within each squirrel species across its geographical distribution using location coordinates (42187 random points) extracted at every 10km distance within IUCN species distribution boundaries.

To investigate the correlation of squirrel coat colour with climate and habitat features, we performed linear mixed-effects models between coat colour and a multivariate combination of the best predictor of each category of environmental variables to investigate coat colour variation under the conflicting conditions of various environmental pressures. We then performed univariate linear mixed-effects models for the overall genus-level data between coat colour and each category of environmental variables to tease apart the individual effects of environmental variables on squirrel coat colouration. For both univariate and multivariate LMM models, species groups and the year of specimen collection were used as random variables to control for changes in fur colouration over time due to long-term storage. Upon inspecting the grouping effects of species as random variables, we replicated the univariate and multivariate analyses for subsets of data within each species to investigate fine-scale species-level responses, with the year of specimen collection as a random variable. Specifically for *F. tristriatus,* we performed multivariate linear mixed-effects models under a partial Bayesian framework due to its low sample size and model complexity.

We performed mean and variance tests to test the difference in the coat colour of squirrel species between raptor and ferret vision. We performed Kruskal-Walis and Dunn tests to test the interspecies variation of the dorsal light and dark stripe dimensions.

With the QCPA framework, we calculated dorsal and lateral colour pattern analysis under three broad categories - colour adjacency analysis (CAA), visual contrast analysis (VCA), and boundary strength analysis (BSA) (Supplementary Table S2) [60,61]. We investigated specific parameters that best describe the stripe pattern characteristics of palm squirrels under each category of analyses. Mean parameters of stripe pattern properties under the colour adjacency, visual contrast, and boundary strength analyses were investigated for their differences across squirrel species, between predator visual systems, and their interactive effects using Aligned Rank-Transform ANOVA [62,63] as a non-parametric equivalent to two-factorial ANOVA.

We performed multivariate regression analyses for each visual system to investigate how the animal’s coat colour and pattern respond to canopy cover and height variation at the genus and species levels. We then performed univariate analysis to tease apart individual effects. For the genus-level data, univariate and multivariate regressions of colour and pattern properties across canopy properties were performed separately for raptor and ferret vision, with species as fixed variables.

We did not perform any explicit phylogenetic methods due to the limited number of species (3) and a lack of phylogenetic information.

## Results

### Coat colour variation across species

*Funambulus* species have four uniquely coloured body regions - body, head, light stripes, and dark stripes. The dark and the light stripes were each species’ most contrasting body regions. The coat (body) colour was typically significantly lighter than the head and darker than the light stripe (Supplementary Tables S3, S4, and Figure S6). Across the three species, *F. pennantii* had the lightest colours for all body regions, while *F*. *tristriatus* had the darkest colours (Supplementary Tables S5, S6, and Figure S7). We found a weak pattern of darkening coat colour associated with an increase in body size for *F. palmarum* and *F. pennantii* and no pattern for *F. tristriatus*. The correlation between body size and coat colour was not significantly different between males and females of each species (Supplementary Figure S8 and Table S7). Upon using latitude and longitude as random variables in linear mixed-effects models, we found no significant differences in coat colour between males and females within each species (Supplementary Table S8).

### Coat colouration seen by predators

The mean coat colour was significantly different, as seen by the ferret and raptor visual systems only for *F. tristriatus*. In contrast, the variance of coat colour was seen significantly better by raptors than ferrets for all three species (Table S9 and Figure S9).

### Coat colour variation across geography

Geographic location significantly determines the coat colour of squirrels. The best GAM model (Adj. R2= 0.509, deviance explained = 55.2%, n=640) includes two-way interactive effects of latitude and longitude with species as a grouping factor (Figure 2 and Supplementary Tables S10, S11). Across the genus, the darkest individuals are predicted along the southwestern coast of India, while the lightest individuals are predicted in the northwestern regions of the Indian peninsula. However, species exhibit different trends across geography.

**Figure 2.**
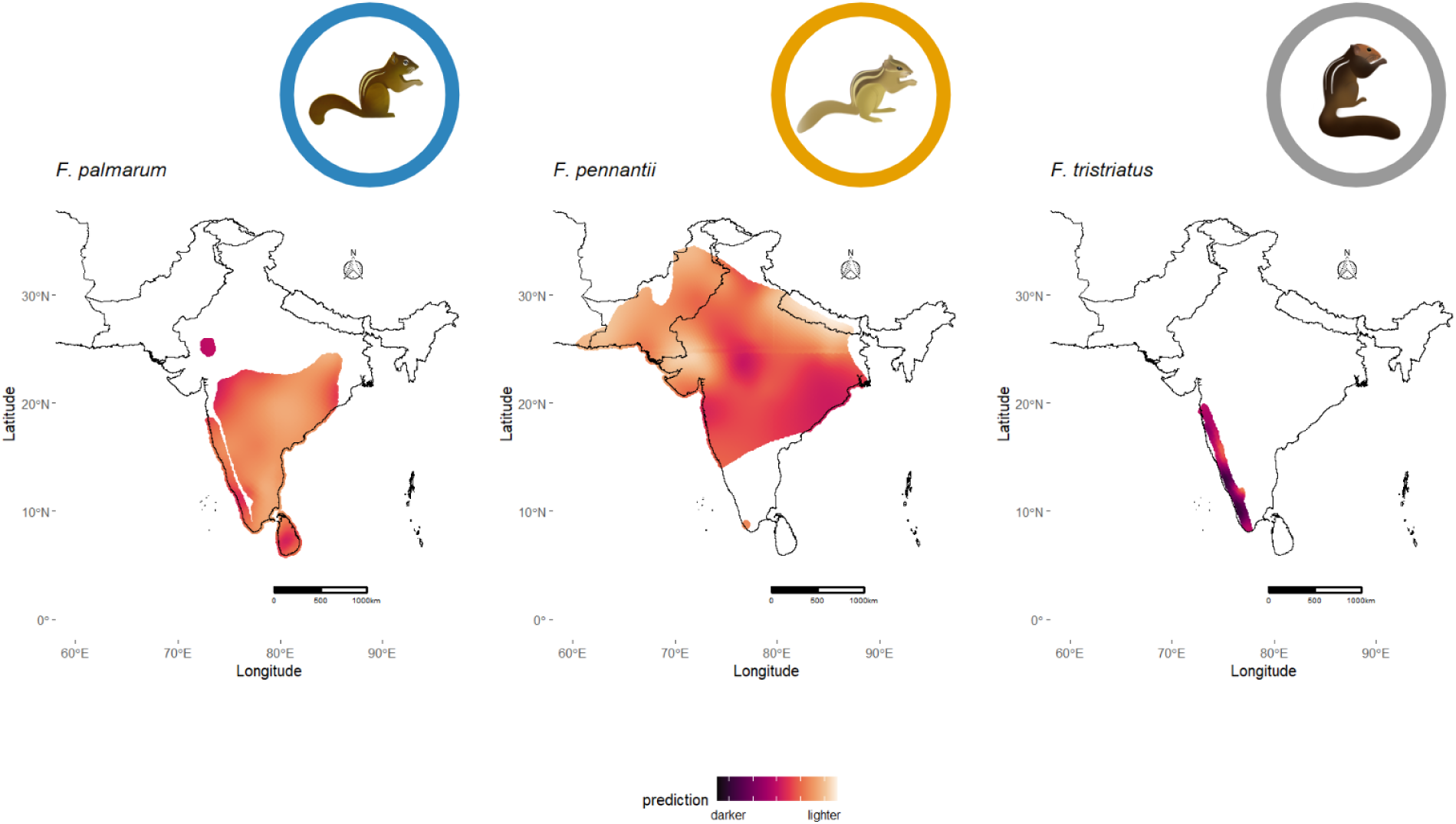
Predicted distribution of coat colour across geographic regions using Generalized additive models. Description: Figure representing the prediction of coat colour throughout the distribution of *Funambulus* species at 42187 points, sampled every 10km. Darker and lighter colour tones represent the locations with the prediction of squirrels having darker and lighter coat colours, respectively. The best model predicts that the darkest individuals (*F. tristratus*) are distributed in southwestern regions, and the lightest individuals (*F. pennantii*) are distributed in north-western regions of the genus range. Each species has a unique axis of lightness-to-darkness transition.

Within *F. palmarum*, the darkest individuals are predicted in the isolated northern Indian population, followed by Sri Lankan individuals, while the lighter individuals are predicted in south-eastern peninsular India (F=1.256, p=5.65E-13). Within *F. pennantii*, the darker individuals are predicted in eastern India, and the lightest individuals are predicted in the northwestern extent of its distribution, including Pakistan and a small region in Iran (F=2.849, p< 2e-16). In *F. tristriatus*, the darkest individuals are predicted in the southern part of the west coast of India, while lighter individuals occur in the northern part of the west coast of India (F=1.58, p=4.59E-10).

### Coat colour associations with the environment

Multivariate mixed-effects models for the genus suggested that precipitation and vegetation (R^2^=0.24) best explain coat colour variation (Supplementary Table S12). Different combinations of environmental variables best describe the coat colour variation for each species. For *F. palmarum,* the best model includes a combination of precipitation and vegetation (R^sq^ =0.24). Vegetation, temperature, and solar radiation (R^sq^=0.16) best explain coat colour variation within *F. pennantii*. Similar to *F. palmarum*, coat colour variation within *F. tristriatus* is best described by the additive terms of precipitation and vegetation (Table 1 and Supplementary Table S12).

**Table 1.** Coat colour associations with environmental variables. Description: Table showing best mixed effects model depicting the association of coat colour with environmental variables for each species and the genus. For the species level data, specimen age was used as a random variable, while for the genus-level data, specimen age, and species categories were used as random variables. After accounting for the confounding effects of multiple environmental variables, precipitation and net primary productivity (vegetation) best determine the coat colour variation of the southern species (*F. palmarum* and *F. tristriatus*) and the genus, while vegetation, temperature, and solar radiation best determine the coat colour variation of the northern species (*F. pennantii*).

|  | *F. palmarum* |  | *F. pennantii* |  | *F. tristriatus* |  | *Genus* |  |
| --- | --- | --- | --- | --- | --- | --- | --- | --- |
| Predictors | Estimates | Std. Error | Estimates | Std. Error | Estimates | Std. Error | Estimates | Std. Error |
| Intercept | -0.07 | 0.13 | -0.65 ** | 0.2 | 1.49 *** | 0.07 | -0.33 | 0.21 |
| average precipitation | -0.39 *** | 0.09 |  |  | -0.09 *** | 0.02 | -0.32 *** | 0.06 |
| net primary productivity | 0.01 | 0.1 | -0.65 *** | 0.17 | -0.18 *** | 0.05 | -0.27 *** | 0.07 |
| average temperature |  |  | -0.45 *** | 0.11 |  |  |  |  |
| solar radiation downwards |  |  | 1.56 *** | 0.44 |  |  |  |  |
| **Random effects** |  |  |  |  |  |  |  |  |
| residual variance | 0.56 |  | 0.56 |  | 0.04 |  | 0.66 |  |
| random intercept variance | 0.10<sub>year</sub> |  | 0.29<sub>year</sub> |  | 0.01<sub>year</sub> |  | 0.17<sub>year</sub> |  |
|  |  |  |  |  |  |  | 0.10<sub>year</sub> |  |
| N | 19<sub>year</sub> |  | 16<sub>year</sub> |  | 15<sub>year</sub> |  | 3<sub>species</sub> |  |
|  |  |  |  |  |  |  | 32<sub>year</sub> |  |
| observations | 174 |  | 287 |  | 64 |  | 525 |  |
| marginal R<sup>2</sup>/conditional R<sup>2</sup> | 0.162/0.291 |  | 0.162/0.445 |  | 0.331/0.412 |  | 0.243/0.463 |  |

Univariate models exhibit similar trends at the genus level, where the coat colour darkens with precipitation and vegetation for all three species. Since species were significantly affected as a random variable, we investigated species-level patterns. In addition to the overall trends, *F. palmarum’s* coat colour darkens with soil organic carbon and lightens with temperature, while the coat colour of *F. tristriatus* significantly lightens with downward solar radiation (Figure 3 and Supplementary Table S13).

**Figure 3.**
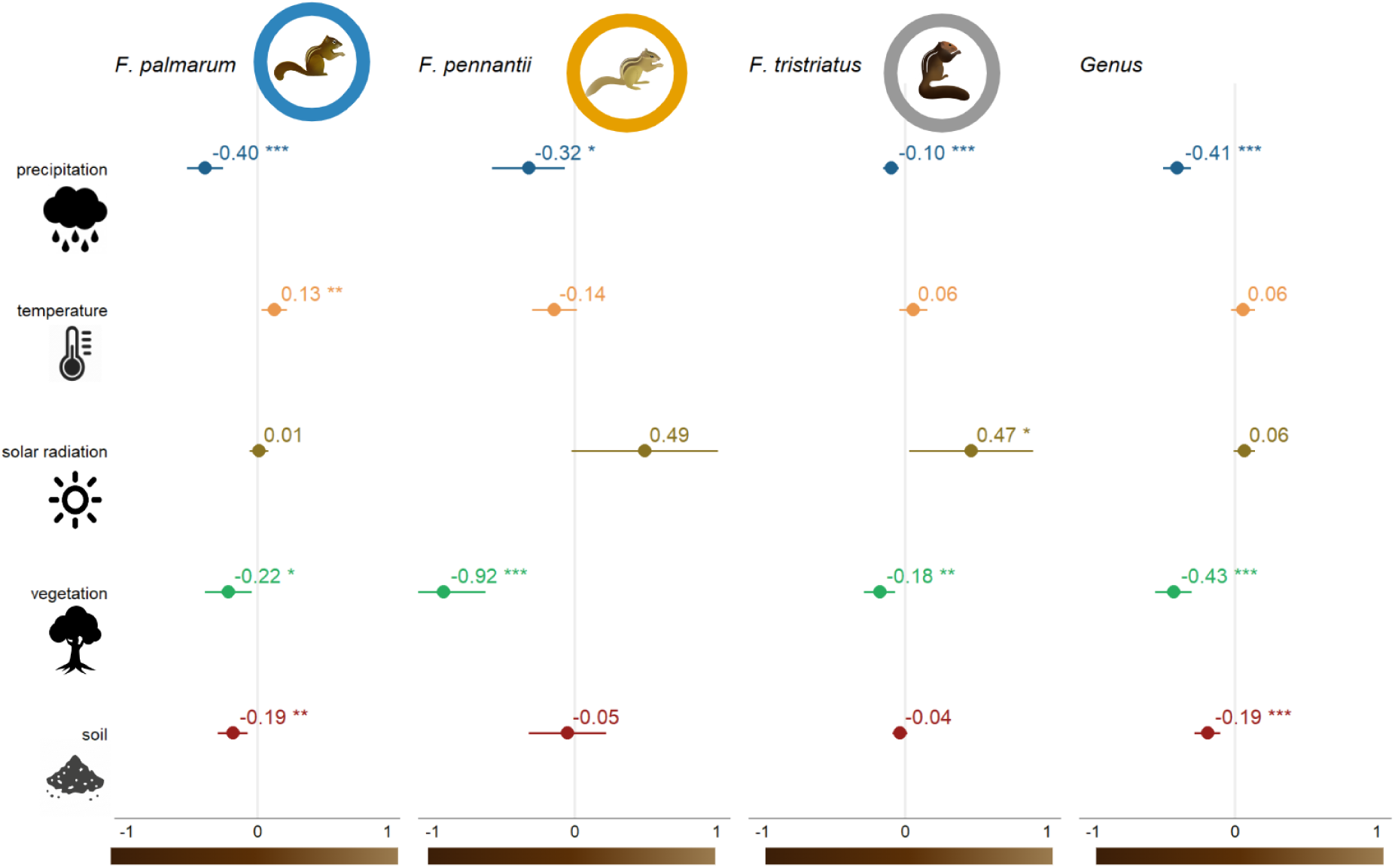
Individual effects of environmental variables on coat colour for each species. Description: Estimate plot depicting the effects of each class of environmental variable on coat colour for individual species. Negative and positive estimates indicate coat colour getting darker and lighter, respectively. For all species, coat colour gets darker with increased precipitation and vegetation. *F. palmarum* shows specific trends with temperature (lighter) and soil colour (darker).

### Interspecific variation of longitudinal stripe dimensions

The three species exhibit significant differences in the length and width dimensions of the first two light and dark stripes. *F. pennantii* had the shortest light and dark stripes, while *F. palmarum* had the longest. Specifically, *F. pennantii* had wider light stripes, similar to *F. palmarum*, and narrower dark stripes. The middle (second) light stripe was narrowest in *F. tristriatus*. Stripe dimensions of *F. palmarum* and *F. tristriatus* have extensive variability and high overlap, while *F. pennantii* has a lower variation (Supplementary Figure S10 and Tables S14, S15).

### Stripes under predator vision

All parameters explaining the dorsal colour adjacency, visual contrast, and boundary strength significantly differed across the three species and the two predator visual systems, with a significant interaction between the squirrel species and visual systems. Overall, raptors can visualize the colour pattern complexity (CAA: Hc, CAA: Ht, and CAA: C), more chromaticity contrast across patches (VCA: MS, VCA: MSsat, and VCA: MDmax), and the colour contrast along boundaries (BSA: MS) better than ferrets can. In contrast, ferrets can visualize more luminance contrast across patches (VCA: MSL and VCA: ML) and saturation (BSA: BMSsat), Dmax (BSA: BMDmax), and luminance contrasts (BSA: BMSL and BSA: BML) along boundaries better than raptors can.

Among the squirrel species, *F. tristriatus* has the highest colour pattern complexity, cluster numbers, and Dmax colour contrast in the raptor vision. *F. pennantii* had the highest luminance contrast across patches (VCA: MSL and VCA: ML) and boundaries (BSA: BMSL and BSA: BML) in the ferret vision. A similar pattern was observed in the lateral patterns of squirrels, with *F. pennantii* having stronger boundaries than the other two squirrel species (Figure 4, Supplementary Table S16 and Supplementary File S2).

**Figure 4.**
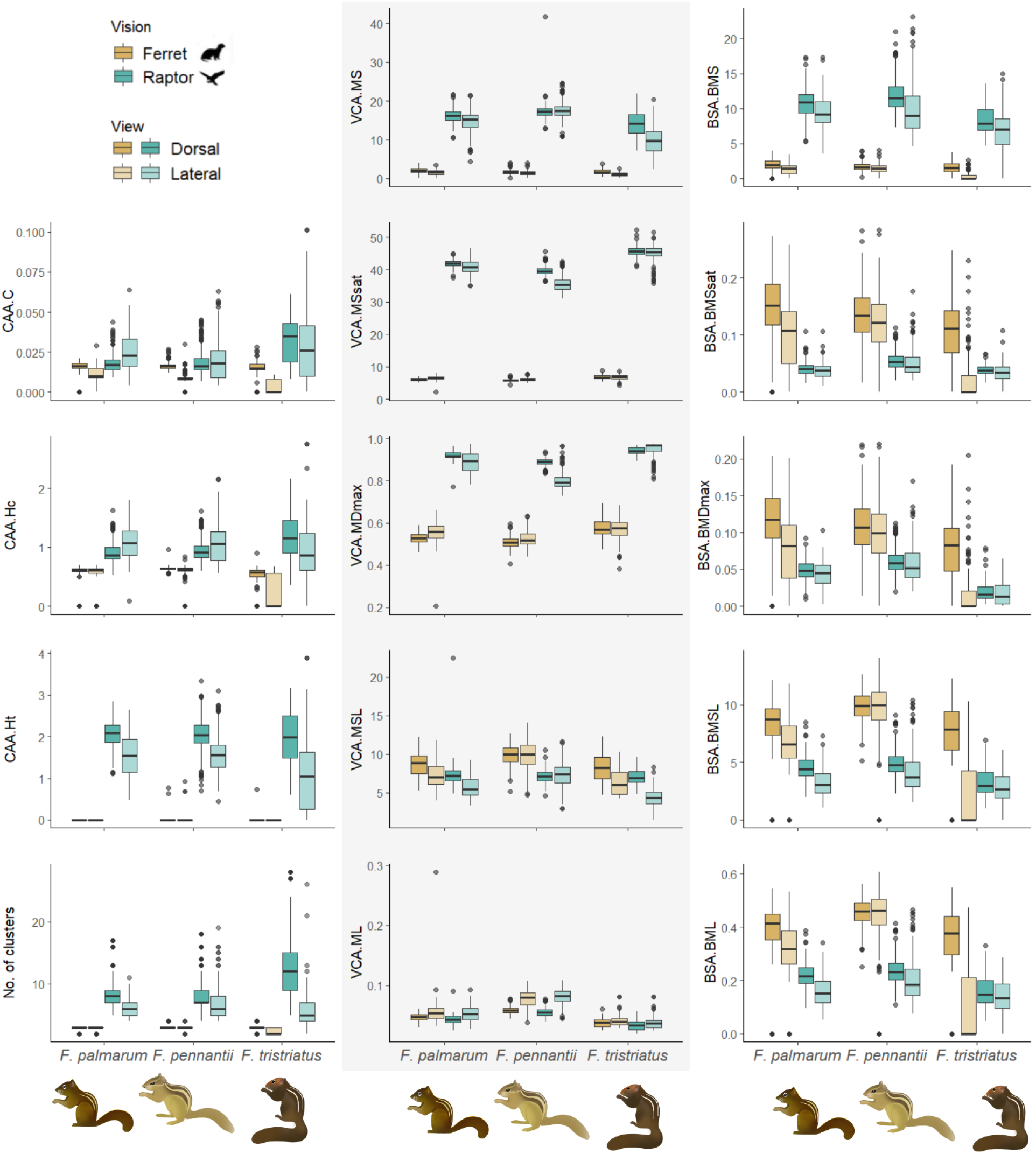
Variation in stripe-pattern parameters under ferret and raptor vision. Description: Boxplots demonstrating the interspecies variation of QCPA parameters: Colour adjacency analysis (A-C), Cluster statistics (D), Visual contrast analysis (E-I), and Boundary strength analysis (J-N). Brown and green boxes indicate ferret and raptor vision, and darker and lighter boxes indicate metrics of dorsal and lateral stripes, respectively. Raptors better visualize the squirrel stripe complexity and visual chromaticity contrasts. Ferrets strongly visualize luminance-based contrasts and boundaries between color-pattern elements. *F. pennantii* has the most contrasting stripes, with stronger boundaries, while *F. tristriatus* has the least contrasting stripes, with weaker boundaries. The lateral stripes of *F. pennantii* have higher contrasts and stronger boundaries equivalent to its dorsal stripes.

### Colour and pattern variation with environmental factors and predator vision

In the raptor and ferret vision, multivariate analyses (MANOVA: Pillai test) of coat colour, VCA.MS and VCA. MSL suggests that *Funambulus* coat colour and pattern contrasts significantly depend on predator species (Raptor: F=28.3432, p< 2.2e-16; Ferret: F=35.235, p< 2.2e-16), tree cover percentage (Raptor: F=11.9743, p=1.34E-07; Ferret: F=13.015, p=3.23E-08), and the interaction between tree cover and forest canopy height (Raptor: F=3.4044, p=0.01752; Ferret: F=5.469, p=0.001043) (Supplementary Table S17, Figures 5 and Supplementary Figure S11).

**Figure 5.**
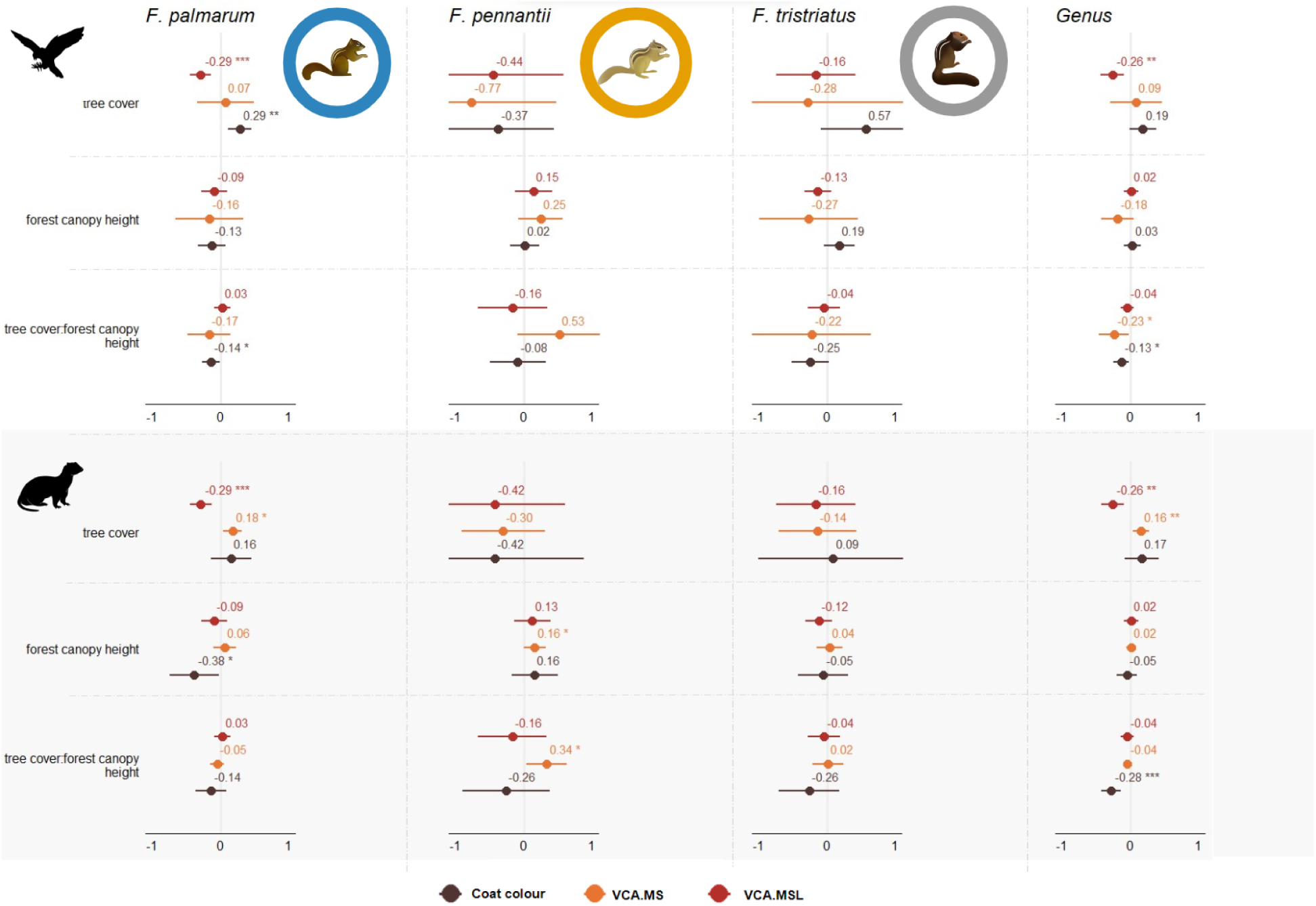
Species-specific patterns of coat colour and stripe contrast variation across canopy properties. Description: Estimate plots displaying the variation in coat colour, stripe colour contrasts, and stripe luminance contrasts with tree cover and forest canopy height. Negative values imply the coat colour getting darker and that the stripe contrasts (colour and luminance) reduce with increasing forest cover and height. For the genus level, the colour of the coat gets darker with increasing tree cover under both ferret and raptor vision. In denser and taller canopies, squirrel colour and luminance contrasts are lower under raptor vision, while only luminance contrast gets lower under ferret vision. Species-specific trends do not show meaningful differences, i.e., colour contrast for raptors and luminance contrasts for ferrets do not vary significantly with forest canopy properties at the species level.

At the species level, *F. palmarum* and *F. pennantii* showed similar variation trends in both the raptor’s and ferret’s vision. Coat colour, VCA.MS and VCA.MSL for *F. palmarum* and *F. pennantii* significantly varies with tree cover and forest canopy height in the ferret vision and only with tree cover in the raptor vision. There was no significant relationship with the canopy properties of *F. tristriatus* in both ferret and raptor vision (Supplementary Table S18 and Figure 5).

In individual models for each colour pattern variable in the raptor vision, coat colour significantly darkens with tree cover (t=-3.035, p=0.00252), with *F. tristriatus* having significantly darker colouration (t=-10.969, p <2e-16). The within-pattern colour contrast (VCA.MS) significantly decreases with an interactive effect between tree cover and forest canopy height (t=-2.104, p=0.0358), suggesting that in cases of increased tree cover, VCA.MS decreases with increasing canopy height and significant differences in squirrel species. We find a similar pattern with luminance contrast (VCA.MSL), suggesting that given increased tree cover, VCA.MSL decreases with increasing canopy height (t=-2.265, p= 0.0239). In the ferret vision, coat colour significantly darkens with an increase in tree cover (t=-3.202, p=0.00144), with *F. tristriatus* having the darkest colouration (t=-10.492, p <2e-16). Colour contrast (VCA.MS) significantly increases with an increase in tree cover (t=2.681,p=0.00757), but no relationship was observed with forest canopy height (t=0.436, p=0.66322) or their interaction term (t=- 1.143, p=0.2534). The within-pattern luminance contrast (VCA.MSL) decreases with an interactive effect between tree cover and forest canopy height (t=-3.964, p=8.38E-05), suggesting that given increased tree cover, VCA.MSL decreases with increasing canopy height (Supplementary Table S19 and Figure 5).

Overall, the model explaining the relationship of VCA.MSL (Raptor: R^2^ = 0.0105, p=0.05625; Ferret: R^2^ =0.2139, p< 2e-16) with canopy properties is poorer in raptor vision and better in ferret vision. In contrast, the model that explains the relationship of VCA.MS (Raptor: VCA.MS: R^2^ =0.167, p< 2e-16; Ferret: R^2^ = 0.07077, p=1.70E-08) with canopy properties is poorer in ferret vision and better in raptor vision. The models explaining body colour’s relationship with canopy properties are stronger for both raptor and ferret vision (Raptor: R^2^ = 0.2754, p< 2e-16; Ferret: R^2^ = 0.2531, p< 2e-16).

Species-specific univariate analysis of coat colour, VCA.MS and VCA.MSL, with tree cover and forest canopy height, also shows different raptor and ferret vision trends. Coat colour significantly decreases with tree cover for the three squirrel species in raptor vision and for *F. palmarum* and *F. tristriatus* in ferret vision. In the raptor vision, only for *F. palmarum,* VCA.MSL significantly increases with tree cover (t=3.352, p=0.00102) but decreases with the interactive term between tree cover and forest canopy height (t=-2.106, p=0.03694). No relationship was found for the other two species. In the ferret vision, VCA.MSL significantly decreases with forest canopy height only for *F. palmarum* (t=-2.104, p=0.0371). VCA.MS significantly increases with tree cover for *F. palmarum* (t=2.594, p=0.0105) and canopy height for *F. pennantii* (t=2.009, p=0.0453). Also, VCA.MS significantly increases with the interaction between tree cover and forest canopy height (t=2.217, p=0.0273). In summary, the relationship between individual colour-pattern properties and canopy properties is weaker at the species level and is relatively more robust at the genus level (Supplementary Table S20 and Figure 5).

## Discussion

Here, we provide evidence that palm squirrel colouration and pattern have evolved in response to multiple abiotic and biotic factors that potentially influence their survival and physiological processes. Stripe patterns with higher contrasts could provide effective motion dazzle effects. A detailed investigation of squirrel stripe properties through the visual systems of their two major predators suggests that the raptor and ferret predators detect stripe contrasts, thereby implying the role of stripes in disruption and motion dazzle. Colour-based stripe contrasts appear more visible to raptors, while luminance-based stripe contrasts appear more visible to ferrets. We also find that squirrels’ overall colour and pattern covary with habitat closure.

### *Funambulus* squirrels show extensive coat colour variation

There is a large variation in the coat colours of *Funambulus* squirrels within and across species. Upon inspecting the patterns of geographic variation of coat colour across the three species of *Funambulus*, we found the darkest individuals across all species along the western coast of southern India, a region with dense evergreen and moist forests. The lightest individuals occurred in the northern latitudes, spreading towards the west, including Pakistan and Iran, with arid to semi-arid habitat types. Several species, including flowers, insects, birds, and mammals, show clinal variations in colouration traits [64–68]. Studies on barn owls [64] and tree squirrels (*Tamiasciurus* spp.) [65] reveal active maintenance of distinct colour phenotypes, even during gene flow between geographically distant closely related species, signifying local adaptations of colouration to environmental pressures.

As expected, we found no significant sexual dichromatism in any *Funambulus* species, suggesting no role of coat colouration in sexual signaling, similar to most non-primate mammalian species [9]. An experimental test of colour vision in *Callosciurus finlaysonii* suggests that the dichromatic vision of tree squirrels cannot differentiate between browns and reds [69], implying the insignificance of colouration in intraspecific signaling.

### Darker squirrels occur in humid and denser habitats

We find that squirrels with darker coats occur in humid and denser habitats, while squirrels with lighter coats occur in dry and open habitats. This pattern is consistent both within and across species. Our results align with several other studies worldwide that establish a darkening of colouration in regions with increased precipitation and vegetation [16,70,71]. The dorsal coat colour of house mice was significantly darker in regions with increased humidity and in closed habitats [72], while that of primates was darker in areas with increased evapotranspiration levels [73]. Darker butterflies in closed habitats suffered lower predation rates than those in open habitats [74], suggesting definite camouflage advantages. Several studies have investigated the pattern of geographical colour variation along Gloger’s rule, which predicts darker colouration in warmer and wetter regions. However, Gloger’s rule does not tease apart the functional advantages for darker individuals living in more humid environments, as the functions could be camouflage-related and physiological [16]. This study tries to find evidence for camouflage and thermal adaptation by teasing apart the effects of environmental variables.

For the genus *Funambulus*, we did not find large-scale patterns that address direct physiological functions of coat colouration since there were no correlates of coat colouration with temperature or solar radiation. However, fine-scale species-level patterns show differential support concerning thermoregulatory and photoprotective functions of coat colouration. In past research, while there is a consistent trend of animals becoming darker in humid habitats [71,73,75–77], there are contradictory patterns for temperature. A few studies support evidence for the thermal melanism hypothesis, where colouration becomes lighter with an increase in temperature [76–78], while others show that colouration darkens with temperature, as suggested by Gloger’s rule [71,77,79,80]. While there were no temperature-related effects in *F. pennantii* and *F. tristriatus*, the coat colour of *F. palmarum* lightens with an increase in temperature. This potentially explains thermal adaptation, where lighter colours absorb lesser heat [81,82], thereby aiding the individuals in regulating body temperature.

We did not find correlates of coat colour with solar radiation for *F. palmarum* and *F. pennantii*. However, in *F. tristriatus*, there was a significant lightening of coat colour with increased solar radiation, contrary to our expectations. An investigation of the evolution of pelage luminance of 151 squirrel species also showed a similar result where dorsal pelage lightens with increasing solar radiation [45]. Given that our metric of shortwave solar was between 200 and 4000uM wavelengths [83], which is not the entire range of potentially harmful radiation (100-400uM) [84], further investigation might be required.

There could also be other variables influencing thermoregulation, such as hair properties like length, diameter, and arrangement or behavioral adaptations [81], which were not considered in our study.

The genus (as well as *F. palmarum)* coat colouration also darkens with increased soil organic carbon content, which typically implies darker soils [45,85], similar to another study on squirrel colouration [45]. *Funambulus* squirrels are scansorial, and they spend a considerable amount of time foraging on the ground [86,87]. Darker coat colour in areas with darker soils implies selection for crypsis via background colour matching.

### Specific stripe elements appear more prominent to specific predators

We found the significance of longitudinal stripes on the bodies of squirrels in anti-predatory functions. Among all three species, the stripe dimensions of *F. pennantii* are relatively conserved in the genus, implying that the trait could be under selection.

Predators visualize stripe contrasts with differential intensities. Visual systems use an interplay of chromatic and achromatic channels [88,89]. Specifically, achromatic information is primarily used for detecting motion [90,91]. This study finds that raptors rely more on colour-based cues, while ferrets rely on luminance-based cues. On investigating the stripe contrast variation across different habitats, we find that the luminance contrasts of the stripes become less pronounced in dense and tall forests and clearer in open habitats with shorter trees. This implies that high-contrast stripes might provide effective motion dazzle effects for both raptors and ferrets in open habitats with shorter trees. Highly contrasting stripes are more conspicuous in open habitats [92]. Additionally, raptors can also see higher colour contrasts in open and short habitats, which might contribute to the motion dazzle effect of the stripes.

Previous studies in snakes [93] and artiodactyls [10] have also established associations between habitat properties and the presence of body patterns, with stripes frequently occurring in open habitats.

### Coat colour and stripes serve specific anti-predatory mechanisms

Colour and pattern co-vary with the environment. Denser forests have darker squirrels with low stripe contrasts, while open habitats have lighter squirrels with high stripe contrasts (Figure 6). Squirrel coat colour could function as a primary anti-predatory mechanism to avoid detection, and the stripes could be a secondary anti-predatory mechanism to avoid attack post-detection.

**Figure 6.**
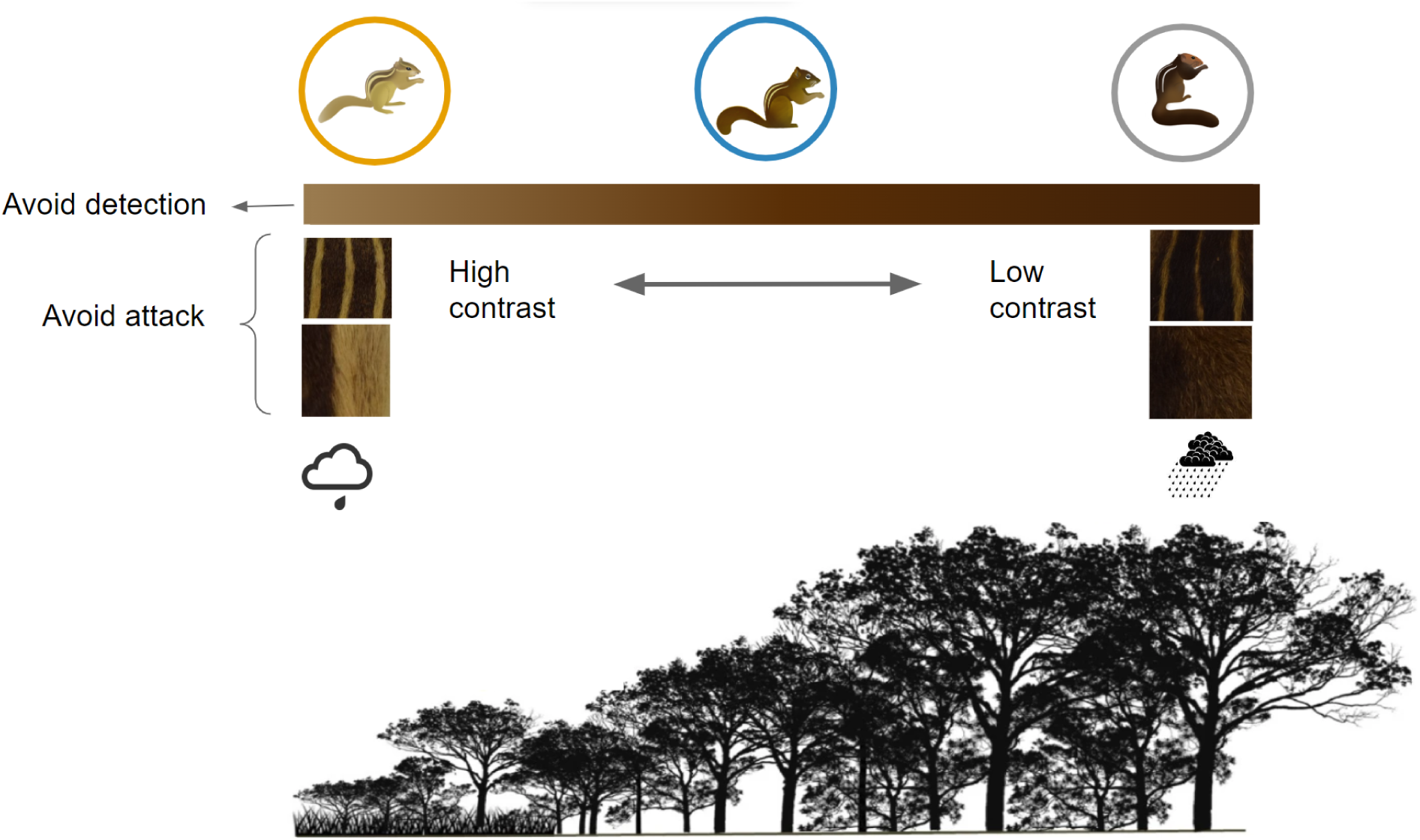
Colour and pattern variation with climate and forest gradient. Description: Squirrel coat colour is darker in more vegetated and humid regions, and stripe-pattern contrast is in forests with denser and taller canopies. Squirrels could use coat colour as a primary anti-predatory mechanism to avoid detection. In open areas, where squirrels are more prone to predator detection, they could gain an advantage with disruptive and motion dazzle effects from highly contrasting stripes. The forest gradient image segment is adapted from [94].

Animals that live in open habitats are more exposed to predators with nowhere to hide, making fleeing and dazzle colouration a significant antipredator defense [93]. Stripes are also associated with group living or sociality in many taxa, including mammals (ruminants) [92], where stripes from multiple individuals produce a stronger signal, thus increasing dazzle camouflage. The contrasting patch sizes also determine the signal’s conspicuousness, where smaller patches are less conspicuous and larger patches are more conspicuous [95].

*F. pennantii,* which live in open habitats with little tree cover [42]*, are the lightest Funambulus and have* the highest pattern contrast. Disruptive colour patterns such as stripes are most effective when the within-pattern contrasts are higher. Our stripe metric analysis also shows that *F. pennantii* has the widest light stripes, making it brighter and more conspicuous. The lighter coat colouration and its strong association with lesser vegetation could aid in crypsis, while highly contrasting parallel stripes could aid their antipredatory behavior once they get detected by the predator. *F. pennantii* also lives in groups of 2-4 individuals [96,97]; hence, investigating the association between group living behaviour and stripe patterns might be interesting.

The conspicuousness of a visual signal for specific predators is sensitive to the light environments in which it is displayed. Structures like closed forests have different light environments from open habitats with little tree cover [90,95,98–100]. Our study modeled the two animal visions under the illumination of the broad daylight spectrum. Understanding the Colour-pattern traits and behavior in mid-understory canopy environments might be valuable.

Studies show that animals often face antagonizing pressures from different selective forces and evolve optimal traits in response [4,101]. This is especially true for multi-functional and complex traits such as colour and patterns across body regions. When facing diverse selective pressures like environmental conditions or predatory risks, an animal achieves optimality through distinct responsive elements on different body regions [102] for different viewers [36,103]. Our study is one of the few to show the evolution of colour and pattern traits concerning the environment, emphasizing their functional differences across two diverse visually hunting predators with strikingly diverse visual capacities.

## Acknowledgments

We thank Abha Gadgil, Aditya Panigrahy, Alina Khan, Divya Janani, Gautami Meherkar, Ranjanee Aron, and Usha Sahoo for their assistance in image processing and pattern quantification; Pooja Gupta for the squirrel illustrations; Rahul Khot and Bombay Natural History Society (BNHS) for access to the squirrel specimens; Swati Udyraj and Aravind PS for image curation; Swati Udayraj, Harsha Kumar and members of the Sciurid Lab for comments on the manuscript. IISER Tirupati and DST-SERB funded this project.

