## Supplementary for "Multifunctional evolution of palm squirrel coat colour and pattern"

### Tables

#### Table S1. Details of environmental variables

| **Environment category** | **Climate/veg parameter** | **Layer** | **Year** | **Original resolution** |
|---|---|---|---|---|
| Geography | Altitude | Global Multi-resolution Terrain Elevation Data 2010 | 2010 | 250m |
| Rainfall & humidity | Rainfall Avg | Chelsa | 02/1979-01/1980 | 1km |
| | Monthly average rainfall | Chelsa | 1978-80 | 1km |
| | Total precipitation | ERA-5 | 1981 | 10km |
| | Relative Humidity | ERA-5 | 1981 | 10km |
| | surface pressure | ERA-5 | 1981 | 10km |
| Temperature | Temperature Avg | Chelsa | 02/1979-01/1980 | 1km |
| | Temperature Min | Chelsa | 02/1979-01/1980 | 1km |
| | Temperature Max | Chelsa | 02/1979-01/1980 | 1km |
| | Monthly average temperature | Chelsa | 1978-80 | 1km |
| | skin temperature | ERA-5 | 1981 | 10km |
| | temperature 2m | ERA-5 | 1981 | 10km |
| | dew point temperature 2m | ERA-5 | 1981 | 10km |
| Solar radiation | surface_net_solar_radiation | ERA-5 | 1981 | 10km |
| | surface_solar_radiation_downwards | ERA-5 | 1981 | 10km |

| | | | | |
|---|---|---|---|---|
| Vegetation | Forest Canopy Height | Global Forest Canopy Height,2019 by Global Land Analysis and Discovery | 2019 | 30m |
| | canopy closure/tree cover | Hansen Global Forest Change v1.8 (2000-2020) | 2000 | ~30m |
| | Landcover | MCD12Q1.006 MODIS Land Cover Type Yearly Global 500m | 2001 | 500m |
| | Net_primary_productivity | MOD17A3HGF.006: Terra Net Primary Production Gap-Filled Yearly Global 500m | 2000 | 500m |
| | NDVI | NOAA CDR AVHRR NDVI: Normalized Difference Vegetation Index, Version 5 | July 1981-June 1982 | 5km |
| Soil | soil texture | Soil grids | 1950-2017 avg | 250m |
| | Soil type | Harmonised World Soil database | 1971-2004 | 1km |
| | soil_bulk_density | OpenLandMap Soil Bulk Density | 2018 | 250 m |
| | soil_organic_carbon_content | OpenLandMap Soil Organic Carbon Content | 2018 | 250 m |
| | E5_soil_temperature_level_1 | ERA-5 | 1981 | 10km |

#### Table S2. Details of stripe-pattern parameters quantified for dorsal and lateral body

| Category | Abbrevation | Parameters |
|---|---|---|
| colour adjacency analysis (n=3) | CAA: C | Shannon colour diversity |
| | CAA: Hc | Shannon transition diversity |
| | CAA: Ht | Pattern complexity |
| Visual contrast analysis (n=5) | VCA: ML | Mean luminance contrast |
| | VCA: MSL | Mean RNL luminance |
| | VCA: MS | Mean colour contrast |
| | VCA: MSL | Mean saturation contrast |
| | VCA: MDmax | Mean Dmax contrast |
| Boundary strength analysis (n=5) | BSA: BML | Mean boundary luminance contrast |
| | BSA: BMSL | Mean boundary RNL luminance |
| | BSA: BMS | Mean boundary colour contrast |
| | BSA: BMSsat | Mean boundary saturation contrast |
| | BSA: BMDmax | Mean boundary Dmax contrast |

#### Table S3. Colour variation across body within species

| Species | | Df | Sum Sq | Mean Sq | F value | Pr(>F) |
|---|---|---|---|---|---|---|
| *F. pennantii* | body region | 4 | 2435.4 | 608.8 | 2736 | <2e-16 *** |
| | residuals | 1812 | 403.2 | 0.2 | | |
| *F. palmarum* | body region | 4 | 1942.4 | 485.6 | 1353 | <2e-16 *** |
| | residuals | 979 | 351.4 | 0.4 | | |
| *F. tristriatus* | body region | 4 | 252.24 | 63.06 | 394.4 | <2e-16 *** |
| | residuals | 385 | 61.56 | 0.16 | | |

#### Table S4. Colour variation between pairs of body regions within species

| Species | Body region | diff | lwr | upr | p adj |
|---|---|---|---|---|---|
| F. pennantii | darkstripe-body | -1.36591514 | -1.46132445 | -1.2705058 | 0 |
| | head-body | -0.96955712 | -1.06522999 | -0.8738842 | 0 |
| | lightstripe-body | 1.85935837 | 1.76394906 | 1.9547677 | 0 |
| | neck-body | -0.88763108 | -0.9830431 | -0.792219 | 0 |
| | head-darkstripe | 0.39635802 | 0.30075043 | 0.4919656 | 0 |

| | | | | | |
|---|---|---|---|---|---|
| | lightstripe-darkstripe | 3.22527351 | 3.12992966 | 3.3206174 | 0 |
| | neck-darkstripe | 0.47668936 | 0.38114818 | 0.5722305 | 0 |
| | lightstripe-head | 2.82891549 | 2.73330789 | 2.9245231 | 0 |
| | neck-head | 0.08033133 | -0.01547305 | 0.1761357 | 0.1486566 |
| | neck-lightstripe | -2.74858415 | -2.84412533 | -2.653043 | 0 |
| F. palmarum | darkstripe-body | -2.0080601 | -2.1730353 | -1.84308482 | 0 |
| | head-body | -1.4045083 | -1.5696939 | -1.23932279 | 0 |
| | lightstripe-body | 1.8502175 | 1.6852423 | 2.01519278 | 0 |
| | neck-body | -1.5334864 | -1.6984617 | -1.36851114 | 0 |
| | head-darkstripe | 0.6035517 | 0.4383662 | 0.76873729 | 0 |
| | lightstripe-darkstripe | 3.8582776 | 3.6933023 | 4.02325285 | 0 |
| | neck-darkstripe | 0.4745737 | 0.3095984 | 0.63954893 | 0 |
| | lightstripe-head | 3.2547259 | 3.0895403 | 3.41991141 | 0 |
| | neck-head | -0.1289781 | -0.2941636 | 0.03620749 | 0.2065418 |
| | neck-lightstripe | -3.3837039 | -3.5486792 | -3.21872867 | 0 |
| F. tristriatus | darkstripe-body | -0.87907848 | -1.0534577 | -0.7046992 | 0 |
| | head-body | -0.598907274 | -0.774999 | -0.4228156 | 0 |
| | lightstripe-body | 1.358845147 | 1.1839079 | 1.5337824 | 0 |
| | neck-body | -0.597168456 | -0.7721057 | -0.4222312 | 0 |
| | head-darkstripe | 0.280171205 | 0.1040795 | 0.4562629 | 0.0001616 |

| | | | | | |
|---|---|---|---|---|---|
| | lightstripe-darkstripe | 2.237923627 | 2.0629864 | 2.4128609 | 0 |
| | neck-darkstripe | 0.281910023 | 0.1069728 | 0.4568473 | 0.0001267 |
| | lightstripe-head | 1.957752421 | 1.7811081 | 2.1343967 | 0 |
| | neck-head | 0.001738818 | -0.1749055 | 0.1783831 | 0.9999999 |
| | neck-lightstripe | -1.956013603 | -2.1315071 | -1.7805201 | 0 |

#### Table S5. Colour variation of a body region across species

| Body region | | Df | Sum Sq | Mean Sq | F value | Pr(>F) |
|---|---|---|---|---|---|---|
| light stripe | Species | 2 | 398.2 | 199.08 | 88.55 | <2e-16 *** |
| | Residuals | 639 | 1436.6 | 2.25 | | |
| dark stripe | Species | 2 | 901.2 | 450.6 | 302.6 | <2e-16 *** |
| | Residuals | 640 | 953 | 1.5 | | |
| body | Species | 2 | 157 | 78.48 | 85.67 | <2e-16 *** |
| | Residuals | 640 | 586.3 | 0.92 | | |
| head | Species | 2 | 630.4 | 315.18 | 168.9 | <2e-16 *** |
| | Residuals | 633 | 1181.3 | 1.87 | | |

Table S6. Pairwise interspecific differences of body color

| Body region | Species pair | diff | lwr | upr | p adj |
|---|---|---|---|---|---|
| lightstripe | *F. pennantii-F. palmarum* | 1.144454 | 0.8333533 | 1.4555546 | 0.00E+00 |
| | *F. tristriatus-F. palmarum* | -1.088415 | -1.559624 | -0.6172064 | 2.00E-07 |
| | *F. tristriatus-F. pennantii* | -2.232869 | -2.6720335 | -1.7937047 | 0.00E+00 |
| darkstripe | *F. pennantii-F. palmarum* | 1.785602 | 1.532408 | 2.038796 | 0 |
| | *F. tristriatus-F. palmarum* | -1.504064 | -1.885822 | -1.122306 | 0 |
| | *F. tristriatus-F. pennantii* | -3.289666 | -3.645216 | -2.934115 | 0 |
| body color | *F. pennantii-F. palmarum* | 0.2073307 | 0.008732111 | 0.4059294 | 0.0383611 |
| | *F. tristriatus-F. palmarum* | -1.3435022 | -1.64294277 | -1.0440615 | 0 |
| | *F. tristriatus-F. pennantii* | -1.5508329 | -1.829716939 | -1.2719488 | 0 |
| head color | *F. pennantii-F. palmarum* | 1.091104 | 0.8067801 | 1.375427 | 0 |
| | *F. tristriatus-F. palmarum* | -1.965385 | -2.3990422 | -1.531727 | 0 |
| | *F. tristriatus-F. pennantii* | -3.056488 | -3.4612189 | -2.651757 | 0 |

#### Table S7. Coat colour variation with sex and body size

| Species | Model fixed effects | Model random effects | AIC | marginal R2 | Std. error | df | F | Pr(>\|t\|) |
|---|---|---|---|---|---|---|---|---|
| *F. pennantii* | body_color | body_size | 808.055 | 0.04868 | 0.912 | 301 | 16.45 | 6.36E-05 |
| *F. pennantii* | body_color | body_size + Sex | 809.6381 | 0.04682 | 0.9129 | 300 | 8.418 | 0.0002774 |
| *F. pennantii* | body_color | Sex | 1015.375 | -0.002654 | 0.9859 | 358 | 0.04958 | 0.8239 |
| *F. pennantii* | body_color | body_size * Sex | 809.5912 | 0.05008 | 0.9113 | 299 | 6.307 | 0.0003679 |
| *F. palamrum* | body_color | body_size | 443.54 | 0.04429 | 0.8883 | 167 | 8.785 | 3.48E-03 |
| *F. palamrum* | body_color | body_size + Sex | 445.3431 | 0.03965 | 0.8904 | 166 | 4.468 | 0.01288 |
| *F. palamrum* | body_color | Sex | 506.2072 | 0.01546 | 0.8888 | 191 | 4.014 | 0.04653 |
| *F. palamrum* | body_color | body_size * Sex | 446.8548 | 0.03662 | 0.8918 | 165 | 3.129 | 0.02729 |
| *F. tristriatus* | body_color | body_size | 24.57516 | 0.05013 | 0.2841 | 57 | 2.557 | 0.08641 |
| *F. tristriatus* | body_color | body_size + Sex | 24.18571 | 0.05013 | 0.2841 | 57 | 2.557 | 0.08641 |
| *F. tristriatus* | body_color | Sex | 17.8207 | 0.01921 | 2.528 | 77 | 2.528 | 0.1159 |
| *F. tristriatus* | body_color | body_size * Sex | 26.17394 | 0.03336 | 0.2866 | 56 | 1.679 | 0.182 |

#### Table S8. Coat colour variation with sex as a fixed variable and geography as a random variable.

| Species | Model fixed effects | Model random effects | Estimate | Std. error | t |
|---|---|---|---|---|---|
| *F. pennantii* | Sex | (1\|Latitude)+(1\|Longitude) | 0.01437 | 0.08875 | 0.162 |

| | | | | | |
|---|---|---|---|---|---|
| *F. palmarum* | Sex | (1\|Latitude)+(1\|Longitude) | -0.03824 | 0.11919 | -0.3 |
| *F. tristriatus* | Sex | (1\|Latitude)+(1\|Longitude) | 0.08175 | 0.04551 | 1.796 |

### Table S9. Differences in colour variation under ferret vs. raptor visual systems

| Test | Species | Test statistic (F/t) | df | p-value | CI lwr | CI high | mean of raptor_f1 | mean of ferret_f1 | Ratio of variances |
|---|---|---|---|---|---|---|---|---|---|
| T-test | *F. palmarum* | 1.003 | 299.41 | 0.3167 | -0.1447485 | 0.4456932 | 0.0177219 | 0.1327505 | |
| | *F. pennantii* | 0.99894 | 591.5 | 0.3182 | -0.1177923 | 0.3616523 | 0.5969871 | 0.4750571 | |
| | *F. tristriatus* | -3.5005 | 122.64 | 0.0006481 | -1.3448783 | -0.3732844 | -2.585904 | -1.758013 | |
| F-test | *F. palmarum* | 2.8361 | 184(num df), 184(denom df) | 4.84E-12 | 2.122445 | 3.789844 | | | 2.836148 |
| | *F. pennantii* | 2.6 | 354(num df), 353(denom df) | < 2.2e-16 | 2.10993 | 3.203701 | | | 2.599961 |
| | *F. tristriatus* | 3.2362 | 78(num df), 76(denom df) | 6.16E-07 | 2.061975 | 5.072456 | | | 3.236233 |

#### Table S10. Best three generalized additive models

| Response variable | Smoother | Model | AIC | ΔAIC | Rsq (adj) | Deviance explained (%) |
|---|---|---|---|---|---|---|
| body color | S | Species+<br>s(Longitude, Latitude, by= Species, k=100, bs="tp",m=1) | **1526.912** | 0 | 0.509 | 55.2 |
| body color | GS | s(Longitude, Latitude, bs="tp",k=100,m=2)+<br>s(Longitude, Latitude, Species, bs="fs", m=2) | 1531.03 | 4.118 | 0.5 | 54.1 |
| body color | GS | s(Longitude, Latitude, bs="tp",k=100,m=2)+<br>s(Longitude, Latitude, Species, bs="fs", m=2)+<br>s(Altitude, Species, bs="fs", m=2) | 1534.816 | 7.904 | 0.499 | 53.9 |

#### Table S11. Smooth terms for the best GAM model

| Smooth terms | edf | Ref. df | F | p-value |
|---|---|---|---|---|
| s(Longitude,Latitude):Species*Funambulus palmarum* | 18.5 | 73 | 1.256 | 5.65E-13 |
| s(Longitude,Latitude):Species*Funambulus pennanti* | 24.89 | 83 | 2.849 | < 2e-16 |
| s(Longitude,Latitude):Species*Funambulus tristriatus* | 11.3 | 40 | 1.58 | 4.59E-10 |

Table S12. Best three mixed effects models

| Data | Response variable | Fixed variables | Random variables | AIC | ΔAIC | Marginal R2 | Conditional R2 |
|---|---|---|---|---|---|---|---|
| Genus | body_color | average_precipitation + net_primary_productivity | (1\|Species)+(1\|year) | 1322.797 | 0 | 0.2429103 | 0.4628197 |
| | body_color | average_precipitation * net_primary_productivity | (1\|Species)+(1\|year) | 1323.616 | 0.819 | 0.2620295 | 0.4735989 |
| | body_color | average_precipitation + solar_radiation_downwards+ net_primary_productivity+ soil_organic_carbon | (1\|Species)+(1\|year) | 1324.79 | 1.993 | 0.2433434 | 0.4632786 |
| *F. palmarum* | body_color | Average_precipitation+ net_primary_productivity | (1\|year) | 418.7623 | 0 | 0.1620623 | 0.2905387 |
| | body_color | average_precipitation* net_primary_productivity | (1\|year) | 418.3222 | -0.4401 | 0.204719 | 0.2956435 |
| | body_color | average_precipitation+ net_primary_productivity+ soil_organic_carbon | (1\|year) | 417.1412 | -1.6211 | 0.2101595 | 0.3195537 |
| *F. pennantii* | body_color | average_temperature + solar_radiation_downwards+ net_primary_productivity | (1\|year) | 685.7736 | 0 | 0.1618168 | 0.445201 |
| | body_color | average_temperature * solar_radiation_downwards* net_primary_productivity | (1\|year) | 685.9412 | 0.1676 | 0.186576 | 0.5127573 |

| | | | | | | | |
|---|---|---|---|---|---|---|---|
| | body_color | average_temperature + solar_radiation_downwards+ net_primary_productivity+ soil_organic_carbon | (1\|year) | 686.8472 | 1.0736 | 0.1650932 | 0.45821 |
| *F. tristriatus* | body_color | average_precipitation+ net_primary_productivity | (1\|year) | -3.01872 | 0 | Bayesian models | |
| | body_color | average_precipitation + average_temperature + net_solar_radiation+ net_primary_productivity+ soil_organic_carbon | (1\|year) | -2.617068 | 0.401652 | | |
| | body_color | average_precipitation+ solar_radiation_downwards+ net_primary_productivity | (1\|year) | -1.014217 | 2.004503 | | |

### Table S13. Mixed effects model with individual environmental variables as fixed variables

| Data | Fixed variables | Random variables | AIC | Marginal R2 | Conditional R2 | Estimate | Std. error | df | t | Pr(>\|t\|) |
|---|---|---|---|---|---|---|---|---|---|---|
| | average_precipitation | (1\|Species)+(1\|year) | 1673.509 | 0.1451033 | 0.3474987 | -0.40838 | 0.04984 | 494.85961 | -8.194 | **2.18E-15** |
| | average_temperature | (1\|Species)+(1\|year) | 1731.225 | 0.002149569 | 0.4433682 | 0.05571 | 0.04251 | 549.81858 | 1.311 | 0.191 |
| | solar_radiation_downwards | (1\|Species)+(1\|year) | 1726.444 | 0.002756475 | 0.445867 | 0.06327 | 0.03885 | 630.83119 | 1.629 | 0.104 |
| | net_primary_productivity | (1\|Species)+(1\|year) | 1351.295 | 0.1398679 | 0.4906502 | -0.43452 | 0.06638 | 450.17189 | -6.546 | **1.62E-10** |
| Genus | soil_organic_carbon | (1\|Species)+(1\|year) | 1692.835 | 0.02762893 | 0.4234038 | -0.19484 | 0.04668 | 574.74768 | -4.174 | **3.46E-05** |
| | average_precipitation | (1\|year) | 968.4051 | 0.02324532 | 0.2093821 | -0.3234 | 0.1297 | 283.2942 | -2.493 | **0.0132** |
| | average_temperature | (1\|year) | 971.34 | 0.01063483 | 0.2276711 | -0.14238 | 0.08003 | 257.19839 | -1.779 | 0.0764 |
| | solar_radiation_downwards | (1\|year) | 965.6872 | 0.01044183 | 0.2350235 | 0.49387 | 0.26189 | 256.57194 | 1.886 | 0.0605 |
| | net_primary_productivity | (1\|year) | 698.6305 | 0.117581 | 0.3551079 | -0.9247 | 0.1521 | 285.435 | -6.078 | **3.89E-09** |

*F. pennantii*

| | | | | | | | | | | |
|---|---|---|---|---|---|---|---|---|---|---|
| | soil_organic_carbon | (1\|year) | 962.7423 | 0.000372562 | 0.199168 | -0.05156 | 0.13805 | 350.09405 | -0.374 | 0.709 |
| *F. palmarum* | average_precipitation | (1\|year) | 470.3236 | 0.1667623 | 0.2811581 | -0.40327 | 0.06928 | 168.37115 | -5.821 | **2.88E-08** |
| | average_temperature | (1\|year) | 492.2659 | 0.04178787 | 0.2702994 | 0.13147 | 0.04911 | 190.09283 | 2.677 | **0.00808** |
| | solar_radiation_downwards | (1\|year) | 498.9944 | 0.0004885183 | 0.2851615 | 0.01208 | 0.03694 | 190.94814 | 0.327 | 0.744 |
| | net_primary_productivity | (1\|year) | 433.9371 | 0.04593045 | 0.3185177 | -0.22451 | 0.09086 | 116.69841 | -2.471 | **0.0149** |
| | soil_organic_carbon | (1\|year) | 481.267 | 0.06944659 | 0.2933041 | -0.18996 | 0.05852 | 161.6597 | -3.246 | **0.00142** |
| *F. tristriatus* | average_precipitation | (1\|year) | 4.67162 | 0.1818039 | 0.4008179 | -0.09768 | 0.02679 | 70.79554 | -3.647 | **0.000504** |
| | average_temperature | (1\|year) | 15.46322 | 0.0184404 | 0.3257921 | 0.05688 | 0.04944 | 69.12291 | 1.15 | 0.254 |
| | solar_radiation_downwards | (1\|year) | 12.39765 | 0.05654351 | 0.3423718 | 0.46743 | 0.21964 | 72.86212 | 2.128 | **0.0367** |
| | net_primary_productivity | (1\|year) | 6.078439 | 0.1692578 | 0.3246279 | -0.17741 | 0.05526 | 25.59246 | -3.21 | **0.00355** |
| | soil_organic_carbon | (1\|year) | 13.60724 | 0.0228755 | 0.3709316 | -0.03673 | 0.02564 | 72.02877 | -1.432 | 0.156 |

### Table S14. Variation in light and dark stripe dimensions across species

| Kruskal-Wallis rank sum test | | | Dunn Test | | | |
|---|---|---|---|---|---|---|
| Chi sq | df | p value | Comparison | Z | P.unadj | P.adj |
| 511.39 | 2 | < 2.2e-16 | F. palmarum - F. pennantii | 21.152678 | 2.61E-99 | 7.82E-99 |
| | | | F. palmarum - F. tristriatus | 2.990245 | 2.79E-03 | 8.36E-03 |
| | | | F. pennantii - F. tristriatus | -13.756738 | 4.64E-43 | 1.39E-42 |

#### Table S15. Variation of individual stripe dimension across species

| Stripe | Dimension | Kruskal-Wallis rank sum test | | | Dunn Test | | | |
|---|---|---|---|---|---|---|---|---|
| | | Chi sq | df | p value | Comparison | Z | P.unadj | P.adj |
| Light stripe 1 | length | 438.89 | 2 | < 2.2e-16 | F. palmarum - F. pennantii | 18.6973737 | 5.20E-78 | 1.56E-77 |
| | | | | | F. palmarum - F. tristriatus | 0.5257391 | 5.99E-01 | 5.99E-01 |
| | | | | | F. pennantii - F. tristriatus | -14.4380487 | 2.98E-47 | 5.96E-47 |
| Light stripe 2 | length | 371.92 | 2 | < 2.2e-16 | F. palmarum - F. pennantii | 17.838471 | 3.55E-71 | 1.07E-70 |
| | | | | | F. palmarum - F. tristriatus | 1.987901 | 4.68E-02 | 4.68E-02 |
| | | | | | F. pennantii - F. tristriatus | -12.159129 | 5.13E-34 | 1.03E-33 |
| Dark stripe 1 | length | 359.9 | 2 | < 2.2e-16 | F. palmarum - F. pennantii | 17.176242 | 4.00E-66 | 1.20E-65 |
| | | | | | F. palmarum - F. tristriatus | 1.034514 | 3.01E-01 | 3.01E-01 |
| | | | | | F. pennantii - F. tristriatus | -12.663884 | 9.37E-37 | 1.87E-36 |
| Dark stripe 2 | length | 399.01 | 2 | < 2.2e-16 | F. palmarum - F. pennantii | 18.195604 | 5.59E-74 | 1.68E-73 |
| | | | | | F. palmarum - F. tristriatus | 1.352106 | 1.76E-01 | 1.76E-01 |

| | | | | | | | | |
|---|---|---|---|---|---|---|---|---|
| | | | | | F. pennantii - F. tristriatus | -13.136955 | 2.02E-39 | 4.04E-39 |
| Light stripe 1 | width | 25.497 | 2 | < 2.2e-16 | F. palmarum - F. pennantii | 0.6701276 | 5.03E-01 | 5.03E-01 |
| | | | | | F. palmarum - F. tristriatus | 4.769379 | 1.85E-06 | 5.54E-06 |
| | | | | | F. pennantii - F. tristriatus | 4.6085848 | 4.05E-06 | 8.11E-06 |
| Light stripe 2 | width | 109.72 | 2 | < 2.2e-16 | F. palmarum - F. pennantii | -0.4151071 | 6.78E-01 | 6.78E-01 |
| | | | | | F. palmarum - F. tristriatus | 9.1549327 | 5.44E-20 | 1.09E-19 |
| | | | | | F. pennantii - F. tristriatus | 10.1534608 | 3.20E-24 | 9.59E-24 |
| Dark stripe 1 | width | 283.41 | 2 | < 2.2e-16 | F. palmarum - F. pennantii | 15.1498547 | 7.59E-52 | 2.28E-51 |
| | | | | | F. palmarum - F. tristriatus | 0.2782225 | 7.81E-01 | 7.81E-01 |
| | | | | | F. pennantii - F. tristriatus | -11.3404502 | 8.27E-30 | 1.65E-29 |
| Dark stripe 2 | width | 425.1 | 2 | < 2.2e-16 | F. palmarum - F. pennantii | 18.5271408 | 1.25E-76 | 3.74E-76 |
| | | | | | F. palmarum - F. tristriatus | 0.2784722 | 7.81E-01 | 7.81E-01 |
| | | | | | F. pennantii - F. tristriatus | -13.9348825 | 3.89E-44 | 7.78E-44 |

### Table S16. Stripe-pattern variation across species and visual systems

| Qcpa parameter | View | Art Anova | | | | | |
|---|---|---|---|---|---|---|---|
| | | | Df | Df.res | F value | Pr(>F) | Sig |
| CAA:C | Dorsal | Species | 2 | 1290 | 63.238 | < 2.22e-16 | *** |
| | | Vision | 1 | 1290 | 141.419 | < 2.22e-16 | *** |
| | | Species:Vision | 2 | 1290 | 154.201 | < 2.22e-16 | *** |
| | Lateral | Species | 2 | 1242 | 28.467 | 8.16E-13 | *** |
| | | Vision | 1 | 1242 | 675.55 | < 2.22e-16 | *** |
| | | Species:Vision | 2 | 1242 | 61.216 | < 2.22e-16 | *** |
| CAA:Hc | Dorsal: | Species | 2 | 1290 | 38.508 | < 2.22e-16 | *** |
| | | Vision | 1 | 1290 | 1567.365 | < 2.22e-16 | *** |
| | | Species:Vision | 2 | 1290 | 124.446 | < 2.22e-16 | *** |
| | Lateral | Species | 2 | 1242 | 54.112 | < 2.22e-16 | |
| | | Vision | 1 | 1242 | 1282.953 | < 2.22e-16 | *** |
| | | Species:Vision | 2 | 1242 | 15.186 | 3.05E-07 | *** |
| CAA:Ht | Dorsal | Species | 2 | 1290 | 52.96 | < 2.22e-16 | *** |
| | | Vision | 1 | 1290 | 6313.034 | < 2.22e-16 | *** |
| | | Species:Vision | 2 | 1290 | 52.887 | < 2.22e-16 | *** |
| | Lateral | Species | 2 | 1242 | 90.994 | < 2.22e-16 | *** |

| | | | | | | | |
|---|---|---|---|---|---|---|---|
| | | Vision | 1 | 1242 | 4130.932 | < 2.22e-16 | *** |
| | | Species:Vision | 2 | 1242 | 87.383 | < 2.22e-16 | *** |
| VCA:MS | Dorsal | Species | 2 | 1274 | 59.219 | < 2.22e-16 | *** |
| | | Vision | 1 | 1274 | 2916.665 | < 2.22e-16 | *** |
| | | Species:Vision | 2 | 1274 | 131.339 | < 2.22e-16 | *** |
| | Lateral | Species | 2 | 1127 | 63.752 | < 2.22e-16 | *** |
| | | Vision | 1 | 1127 | 1583.253 | < 2.22e-16 | *** |
| | | Species:Vision | 2 | 1127 | 228.618 | < 2.22e-16 | *** |
| VCA:MSsat | Dorsal | Species | 2 | 1274 | 1148.17 | < 2.22e-16 | *** |
| | | Vision | 1 | 1274 | 2924.07 | < 2.22e-16 | *** |
| | | Species:Vision | 2 | 1274 | 479.06 | < 2.22e-16 | *** |
| | Lateral | Species | 2 | 1127 | 913.32 | < 2.22e-16 | *** |
| | | Vision | 1 | 1127 | 1613.61 | < 2.22e-16 | *** |
| | | Species:Vision | 2 | 1127 | 303.05 | < 2.22e-16 | *** |
| VCA:MDmaX | Dorsal | Species | 2 | 1274 | 613.832 | < 2.22e-16 | *** |
| | | Vision | 1 | 1274 | 2896.386 | < 2.22e-16 | *** |
| | | Species:Vision | 2 | 1274 | 23.698 | 7.85E-11 | *** |
| | Lateral | Species | 2 | 1127 | 671.006 | < 2.22e-16 | *** |
| | | Vision | 1 | 1127 | 1601.572 | < 2.22e-16 | *** |
| | | Species:Vision | 2 | 1127 | 81.753 | < 2.22e-16 | *** |
| VCA:MSL | Dorsal | Species | 2 | 1274 | 57.961 | < 2.22e-16 | *** |
| | | Vision | 1 | 1274 | 848.339 | < 2.22e-16 | *** |
| | | Species:Vision | 2 | 1274 | 64.469 | < 2.22e-16 | *** |
| | Lateral | Species | 2 | 1127 | 365.566 | < 2.22e-16 | *** |
| | | Vision | 1 | 1127 | 390.875 | < 2.22e-16 | *** |

| | | | | | | | |
|---|---|---|---|---|---|---|---|
| | | Species:Vision | 2 | 1127 | 10.722 | 2.44E-05 | *** |
| VCA:ML | Dorsal | Species | 2 | 1274 | 1010.88 | < 2.22e-16 | *** |
| | | Vision | 1 | 1274 | 91.878 | < 2.22e-16 | *** |
| | | Species:Vision | 2 | 1274 | 2.114 | 0.12117 | |
| | Lateral | Species | 2 | 1127 | 623.2309 | < 2.22e-16 | *** |
| | | Vision | 1 | 1127 | 9.6303 | 0.0019615 | ** |
| | | Species:Vision | 2 | 1127 | 11.7049 | 9.30E-06 | *** |
| BSA: BMS | Dorsal | Species | 2 | 1290 | 120.25 | < 2.22e-16 | *** |
| | | Vision | 1 | 1290 | 3007.53 | < 2.22e-16 | *** |
| | | Species:Vision | 2 | 1290 | 118.99 | < 2.22e-16 | *** |
| | Lateral | Species | 2 | 1242 | 93.905 | < 2.22e-16 | *** |
| | | Vision | 1 | 1242 | 2955.552 | < 2.22e-16 | *** |
| | | Species:Vision | 2 | 1242 | 26.784 | 4.09E-12 | *** |
| BSA: BMSsat | Dorsal | Species | 2 | 1290 | 55.72 | < 2.22e-16 | *** |
| | | Vision | 1 | 1290 | 1503.134 | < 2.22e-16 | *** |
| | | Species:Vision | 2 | 1290 | 47.266 | < 2.22e-16 | *** |
| | Lateral | Species | 2 | 1242 | 192.87 | < 2.22e-16 | *** |
| | | Vision | 1 | 1242 | 511.19 | < 2.22e-16 | *** |
| | | Species:Vision | 2 | 1242 | 122.68 | < 2.22e-16 | *** |
| BSA: BMDmax | Dorsal | Species | 2 | 1290 | 128.327 | < 2.22e-16 | *** |
| | | Vision | 1 | 1290 | 1009.638 | < 2.22e-16 | *** |
| | | Species:Vision | 2 | 1290 | 22.678 | 2.09E-10 | *** |
| | Lateral | Species | 2 | 1242 | 241.921 | < 2.22e-16 | *** |
| | | Vision | 1 | 1242 | 232.643 | < 2.22e-16 | *** |
| | | Species:Vision | 2 | 1242 | 51.675 | < 2.22e-16 | *** |

| | | | | | | | |
|---|---|---|---|---|---|---|---|
| BSA: BMSL | Dorsal | Species | 2 | 1290 | 153.581 | < 2.22e-16 | *** |
| | | Vision | 1 | 1290 | 1902.825 | < 2.22e-16 | *** |
| | | Species:Vision | 2 | 1290 | 17.793 | 2.38E-08 | *** |
| | Lateral | Species | 2 | 1242 | 420.21 | < 2.22e-16 | *** |
| | | Vision | 1 | 1242 | 809.19 | < 2.22e-16 | *** |
| | | Species:Vision | 2 | 1242 | 232.65 | < 2.22e-16 | *** |
| BSA: BML | Dorsal | Species | 2 | 1290 | 151.326 | < 2.22e-16 | *** |
| | | Vision | 1 | 1290 | 1967.572 | < 2.22e-16 | *** |
| | | Species:Vision | 2 | 1290 | 14.204 | 7.91E-07 | *** |
| | Lateral | Species | 2 | 1242 | 400.88 | < 2.22e-16 | *** |
| | | Vision | 1 | 1242 | 811.49 | < 2.22e-16 | *** |
| | | Species:Vision | 2 | 1242 | 225.46 | < 2.22e-16 | *** |

Table S17. Multivariate colour and pattern variation across canopy closure and height (Type 3 Manova)

| Vision | Variable | Df | test statistic | approx F | num DF | den DF | Pr(>]t]) |
|---|---|---|---|---|---|---|---|
| Raptor | Species | 2 | 0.271647 | 28.3432 | 6 | 1082 | < 2.2e-16 |
| | tree cover perc | 1 | 0.062375 | 11.9743 | 3 | 540 | 1.34E-07 |
| | forest canopy height | 1 | 0.005725 | 1.0364 | 3 | 540 | 0.37598 |
| | tree_cover_perc:forest_canopy_height | 1 | 0.018562 | 3.4044 | 3 | 540 | 0.01752 |
| Ferret | Species | 2 | 0.32842 | 35.235 | 6 | 1076 | < 2.2e-16 |
| | tree cover perc | 1 | 0.06778 | 13.015 | 3 | 537 | 3.23E-08 |
| | forest canopy height | 1 | 0.00285 | 0.512 | 3 | 537 | 0.673937 |
| | tree_cover_perc:forest_canopy_height | 1 | 0.02965 | 5.469 | 3 | 537 | 0.001043 |

Table S18. Species-specific multivariate colour and pattern variation across canopy closure and height (Type 3 Manova)

| Vision | Species | Variable | Df | test statistic | approx F | num DF | den DF | Pr(>]t]) |
|---|---|---|---|---|---|---|---|---|
| Raptor | *F. palmarum* | tree cover perc | 1 | 0.162771 | 9.2672 | 3 | 143 | 1.22E-05 |
| | | forest canopy height | 1 | 0.051054 | 2.5645 | 3 | 143 | 0.05705 |
| | | tree_cover_perc:forest_canopy_height | 1 | 0.035842 | 1.772 | 3 | 143 | 0.1552 |
| | *F. pennantii* | tree cover perc | 1 | 0.035603 | 3.9379 | 3 | 320 | 0.008809 |
| | | forest canopy height | 1 | 0.019702 | 2.1438 | 3 | 320 | 0.094618 |
| | | tree_cover_perc:forest_canopy_height | 1 | 0.010603 | 1.1432 | 3 | 320 | 0.331747 |
| | *F. tristriatus* | tree cover perc | 1 | 0.067413 | 1.518 | 3 | 63 | 0.2185 |
| | | forest canopy height | 1 | 0.066359 | 1.4926 | 3 | 63 | 0.2252 |
| | | tree_cover_perc:forest_canopy_height | 1 | 0.064326 | 1.4437 | 3 | 63 | 0.2385 |
| Ferret | *F. palmarum* | tree cover perc | 1 | 0.159934 | 8.948 | 3 | 141 | 1.82E-05 |
| | | forest canopy height | 1 | 0.06582 | 3.3115 | 3 | 141 | 0.02195 |
| | | tree_cover_perc:forest_canopy_height | 1 | 0.018867 | 0.9038 | 3 | 141 | 0.44106 |
| | *F. pennantii* | tree cover perc | 1 | 0.034212 | 3.8612 | 3 | 327 | 0.009746 |
| | | forest canopy height | 1 | 0.032876 | 3.7053 | 3 | 327 | 0.012011 |
| | | tree_cover_perc:forest_canopy_height | 1 | 0.017081 | 1.8941 | 3 | 327 | 0.130375 |
| | *F. tristriatus* | tree cover perc | 1 | 0.111128 | 2.41708 | 3 | 58 | 0.07547 |
| | | forest canopy height | 1 | 0.027602 | 0.54878 | 3 | 58 | 0.65097 |
| | | tree_cover_perc:forest_canopy_height | 1 | 0.026378 | 0.5238 | 3 | 58 | 0.66764 |

### Table S19. Coat colour and stripe contrast variation across canopy closure and height

| Vision | Response variable | Independent variable | Estimate | Std. Error | t value | Pr(>]t]) | Adj Rsq | F statistic | DF | p-value |
|---|---|---|---|---|---|---|---|---|---|---|
| | | (Intercept) | 0.15153 | 0.08484 | 1.786 | 0.07465 | 0.2754 | 42.57 | 5, 542 | < 2e-16 |
| | | SpeciesF. pennantii | -0.1369 | 0.11086 | -1.235 | 0.2174 | | | | |
| | | SpeciesF. tristriatus | -1.53157 | 0.13963 | -10.969 | <2e-16 | | | | |
| | | tree cover perc | -0.2593 | 0.08545 | -3.035 | 0.00252 | | | | |
| | | forest_canopy_height | 0.01624 | 0.05282 | 0.307 | 0.7586 | | | | |
| | body_color | tree cover: forest canopy height | -0.04456 | 0.04911 | -0.907 | 0.36457 | | | | |
| | | (Intercept) | 16.65355 | 0.19231 | 86.597 | <2e-16 | 0.167 | 22.93 | 5, 542 | < 2e-16 |
| | | SpeciesF. pennantii | 0.55303 | 0.25129 | 2.201 | 0.0282 | | | | |
| | | SpeciesF. tristriatus | -1.76824 | 0.31651 | -5.587 | 3.67E-08 | | | | |
| | | tree cover perc | 0.08584 | 0.19368 | 0.443 | 0.6578 | | | | |
| | | forest_canopy_height | -0.18104 | 0.11973 | -1.512 | 0.1311 | | | | |
| Raptor | VCA.MS | tree cover: forest canopy height | -0.23424 | 0.11131 | -2.104 | 0.0358 | | | | |

| | | | | | | | | | | |
|---|---|---|---|---|---|---|---|---|---|---|
| | VCA.MSL | (Intercept) | 7.36066 | 0.09806 | 75.065 | <2e-16 | 0.01057 | 2.168 | 5, 542 | 0.05625 |
| | | SpeciesF. pennantii | -0.13304 | 0.12813 | -1.038 | 0.2996 | | | | |
| | | SpeciesF. tristriatus | -0.08135 | 0.16138 | -0.504 | 0.6144 | | | | |
| | | tree cover perc | 0.18725 | 0.09876 | 1.896 | 0.0585 | | | | |
| | | forest_canopy_height | 0.03318 | 0.06105 | 0.544 | 0.587 | | | | |
| | | tree cover: forest canopy height | -0.12857 | 0.05676 | -2.265 | 0.0239 | | | | |
| | body_color | (Intercept) | 0.15824 | 0.08514 | 1.859 | 0.06364 | 0.2531 | 37.87 | 5, 539 | <2e-16 |
| | | SpeciesF. pennantii | -0.15248 | 0.11058 | -1.379 | 0.16851 | | | | |
| | | SpeciesF. tristriatus | -1.49319 | 0.14231 | -10.492 | <2e-16 | | | | |
| | | tree cover perc | -0.27314 | 0.0853 | -3.202 | 0.00144 | | | | |
| | | forest_canopy_height | 0.02493 | 0.05261 | 0.474 | 0.63583 | | | | |
| | | tree cover: forest canopy height | -0.03402 | 0.04869 | -0.699 | 0.48502 | | | | |
| Ferret | VCA.MS | (Intercept) | 1.93951 | 0.05952 | 32.589 | <2e-16 | 0.07077 | 9.286 | 5, 539 | 1.70E-08 |
| | | SpeciesF. pennantii | -0.18871 | 0.0773 | -2.441 | 0.01495 | | | | |
| | | SpeciesF. tristriatus | -0.26256 | 0.09948 | -2.639 | 0.00854 | | | | |
| | | tree cover perc | 0.15982 | 0.05962 | 2.681 | 0.00757 | | | | |
| | | forest_canopy_height | 0.01602 | 0.03678 | 0.436 | 0.66322 | | | | |

| | | | | | | | | | | |
|---|---|---|---|---|---|---|---|---|---|---|
| | | tree cover: forest canopy height | -0.03892 | 0.03404 | -1.143 | 0.2534 | | | | |
| | | (Intercept) | 8.94841 | 0.12505 | 71.559 | <2e-16 | 0.2139 | 30.61 | 5, 539 | <2e-16 |
| | | SpeciesF. pennantii | 1.07472 | 0.16241 | 6.617 | 8.84E-11 | | | | |
| | | SpeciesF. tristriatus | -0.18286 | 0.20901 | -0.875 | 0.382 | | | | |
| | | tree cover perc | 0.16884 | 0.12528 | 1.348 | 0.178 | | | | |
| | | forest_canopy_height | -0.04783 | 0.07727 | -0.619 | 0.536 | | | | |
| | VCA.MSL | tree cover: forest canopy height | -0.28346 | 0.07152 | -3.964 | 8.38E-05 | | | | |

Table S20. Species-specific patterns of coat colour and stripe contrast variation across canopy closure and height

| Vision | Species | Dependent variable | Independent variable | Estimate | Std. Error | t value | Pr(>]t]) | Adj Rsq | F statistic | DF | p-value |
|---|---|---|---|---|---|---|---|---|---|---|---|
| Raptor | *F. palmarum* | body_color | (Intercept) | 0.14203 | 0.07794 | 1.822 | 0.070488 | 0.2069 | 13.87 | 3, 145 | 5.34E-08 |
| | | | tree cover perc | -0.29031 | 0.07957 | -3.649 | 0.000367 | | | | |
| | | | forest_canopy_height | -0.09152 | 0.09562 | -0.957 | 0.340098 | | | | |
| | | | tree cover: forest canopy height | 0.03206 | 0.06025 | 0.532 | 0.595462 | | | | |
| | | VCA.MS | (Intercept) | 16.55688 | 0.20604 | 80.357 | <2e-16 | 0.0655 | 1.83 | 3, 145 | 0.1442 |

| | | | | | | | | | | | |
|---|---|---|---|---|---|---|---|---|---|---|---|
| | | | tree cover perc | 0.07323 | 0.21034 | 0.348 | 0.728 | | | | |
| | | | forest_canopy_ height | -0.16023 | 0.25278 | -0.634 | 0.527 | | | | |
| | | | tree cover: forest canopy height | -0.16524 | 0.15928 | -1.037 | 0.301 | | | | |
| | | VCA.MSL | (Intercept) | 7.28353 | 0.08336 | 87.374 | <2e-16 | 6.15E-02 | 4.232 | 3, 145 | 0.006689 |
| | | | tree cover perc | 0.28529 | 0.0851 | 3.352 | 0.00102 | | | | |
| | | | forest_canopy_ height | -0.12627 | 0.10227 | -1.235 | 0.21895 | | | | |
| | | | tree cover: forest canopy height | -0.1357 | 0.06444 | -2.106 | 0.03694 | | | | |
| | *F. pennantii* | body_color | (Intercept) | 0.02032 | 0.2201 | 0.092 | 0.927 | 0.02025 | 3.239 | 3, 322 | 0.02242 |
| | | | tree cover perc | -0.43955 | 0.52182 | -0.842 | 0.4 | | | | |
| | | | forest_canopy_ height | 0.14529 | 0.13716 | 1.059 | 0.29 | | | | |
| | | | tree cover: forest canopy height | -0.16479 | 0.25874 | -0.637 | 0.525 | | | | |
| | | VCA.MS | (Intercept) | 16.7935 | 0.267 | 62.893 | <2e-16 | 0.0008209 | 1.089 | 3, 322 | 0.3539 |
| | | | tree cover perc | -0.7653 | 0.633 | -1.209 | 0.2276 | | | | |
| | | | forest_canopy_ height | 0.2513 | 0.1664 | 1.51 | 0.1319 | | | | |
| | | | tree cover: forest canopy height | 0.529 | 0.3139 | 1.685 | 0.0929 | | | | |

| | | | | | | | | | | | |
|---|---|---|---|---|---|---|---|---|---|---|---|
| | | VCA.MSL | (Intercept) | 6.98318 | 0.17487 | 39.934 | <2e-16 | 0.008112 | 1.886 | 3, 322 | 0.1318 |
| | | | tree cover perc | -0.37444 | 0.41458 | -0.903 | 0.367 | | | | |
| | | | forest_canopy_height | 0.01991 | 0.10897 | 0.183 | 0.855 | | | | |
| | | | tree cover: forest canopy height | -0.08392 | 0.20556 | -0.408 | 0.683 | | | | |
| | *F. tristriatus* | body_color | (Intercept) | -1.36567 | 0.13951 | -9.789 | 2.07E-14 | 0.1512 | 5.036 | 3, 65 | 0.003363 |
| | | | tree cover perc | -0.15847 | 0.28969 | -0.547 | 0.586 | | | | |
| | | | forest_canopy_height | -0.1346 | 0.09706 | -1.387 | 0.17 | | | | |
| | | | tree cover: forest canopy height | -0.03848 | 0.11791 | -0.326 | 0.745 | | | | |
| | | VCA.MS | (Intercept) | 14.9633 | 0.5163 | 28.979 | <2e-16 | 0.06799 | 2.653 | 3, 65 | 0.05588 |
| | | | tree cover perc | -0.2844 | 1.0722 | -0.265 | 0.792 | | | | |
| | | | forest_canopy_height | -0.2671 | 0.3592 | -0.743 | 0.46 | | | | |
| | | | tree cover: forest canopy height | -0.2218 | 0.4364 | -0.508 | 0.613 | | | | |
| | | VCA.MSL | (Intercept) | 7.2316 | 0.1601 | 45.163 | <2e-16 | 0.1113 | 3.839 | 3, 65 | 0.01358 |
| | | | tree cover perc | 0.5744 | 0.3325 | 1.728 | 0.0888 | | | | |
| | | | forest_canopy_height | 0.1858 | 0.1114 | 1.668 | 0.1002 | | | | |

| | | | | | | | | | | | |
|---|---|---|---|---|---|---|---|---|---|---|---|
| | | | tree cover: forest canopy height | -0.2452 | 0.1353 | -1.812 | 0.0746 | | | | |
| Ferret | *F. palmarum* | body_color | (Intercept) | 0.14087 | 0.07607 | 1.852 | 0.066126 | 0.2126 | 14.14 | 3, 143 | 4.03E-08 |
| | | | tree cover perc | -0.29787 | 0.07784 | -3.827 | 0.000194 | | | | |
| | | | forest_canopy_height | -0.0712 | 0.09444 | -0.754 | 0.452148 | | | | |
| | | | tree cover: forest canopy height | 0.03398 | 0.05839 | 0.582 | 0.561463 | | | | |
| | | VCA.MS | (Intercept) | 1.90841 | 0.06758 | 28.238 | <2e-16 | 0.0765 | 5.031 | 3, 143 | 0.00241 |
| | | | tree cover perc | 0.17941 | 0.06915 | 2.594 | 0.0105 | | | | |
| | | | forest_canopy_height | 0.06351 | 0.0839 | 0.757 | 0.4503 | | | | |
| | | | tree cover: forest canopy height | -0.04776 | 0.05187 | -0.921 | 0.3588 | | | | |
| | | VCA.MSL | (Intercept) | 8.9489 | 0.1457 | 61.434 | <2e-16 | 0.07748 | 5.088 | 3, 143 | 0.002242 |
| | | | tree cover perc | 0.1607 | 0.1491 | 1.078 | 0.2827 | | | | |
| | | | forest_canopy_height | -0.3805 | 0.1808 | -2.104 | 0.0371 | | | | |
| | | | tree cover: forest canopy height | -0.1382 | 0.1118 | -1.236 | 0.2185 | | | | |
| | *F. pennantii* | body_color | (Intercept) | 0.01563 | 0.2193 | 0.071 | 0.943 | 0.01797 | 3.026 | 3, 329 | 0.02973 |
| | | | tree cover perc | -0.41534 | 0.51969 | -0.799 | 0.425 | | | | |

| | | | | | | | | | | | |
|---|---|---|---|---|---|---|---|---|---|---|---|
| | | | forest_canopy_ height | 0.12907 | 0.13487 | 0.957 | 0.339 | | | | |
| | | | tree cover: forest canopy height | -0.16401 | 0.25568 | -0.641 | 0.522 | | | | |
| | | VCA.MS | (Intercept) | 1.54894 | 0.13065 | 11.856 | <2e-16 | 0.01841 | 3.075 | 3, 329 | 0.02783 |
| | | | tree cover perc | -0.29752 | 0.3096 | -0.961 | 0.3373 | | | | |
| | | | forest_canopy_ height | 0.16146 | 0.08035 | 2.009 | 0.0453 | | | | |
| | | | tree cover: forest canopy height | 0.33768 | 0.15232 | 2.217 | 0.0273 | | | | |
| | | VCA.MSL | (Intercept) | 9.8627 | 0.2795 | 35.29 | <2e-16 | 0.01952 | 3.204 | 3, 329 | 0.02346 |
| | | | tree cover perc | -0.4244 | 0.6623 | -0.641 | 0.522 | | | | |
| | | | forest_canopy_ height | 0.1614 | 0.1719 | 0.939 | 0.348 | | | | |
| | | | tree cover: forest canopy height | -0.2581 | 0.3258 | -0.792 | 0.429 | | | | |
| | *F. tristriatus* | body_color | (Intercept) | -1.32447 | 0.13943 | -9.499 | 1.44E-13 | 0.1286 | 4.099 | 3, 60 | 0.01032 |
| | | | tree cover perc | -0.15763 | 0.28852 | -0.546 | 0.587 | | | | |
| | | | forest_canopy_ height | -0.12526 | 0.09755 | -1.284 | 0.204 | | | | |
| | | | tree cover: forest canopy height | -0.03157 | 0.1162 | -0.272 | 0.787 | | | | |
| | | VCA.MS | (Intercept) | 1.64677 | 0.13696 | 12.023 | <2e-16 | -0.0371 | 0.2487 | 3, 60 | 0.8619 |
| | | | tree cover perc | -0.14016 | 0.28341 | -0.495 | 0.623 | | | | |

| | | | | | | | | | | | |
|---|---|---|---|---|---|---|---|---|---|---|---|
| | | | forest_canopy_ height | 0.03699 | 0.09582 | 0.386 | 0.701 | | | | |
| | | | tree cover: forest canopy height | 0.0151 | 0.11415 | 0.132 | 0.895 | | | | |
| | | VCA.MSL | (Intercept) | 8.7283 | 0.26383 | 33.083 | <2e-16 | 0.07586 | 2.724 | 3, 60 | 0.05209 |
| | | | tree cover perc | 0.09357 | 0.54594 | 0.171 | 0.864 | | | | |
| | | | forest_canopy_ height | -0.05242 | 0.18458 | -0.284 | 0.777 | | | | |
| | | | tree cover: forest canopy height | -0.26097 | 0.21988 | -1.187 | 0.24 | | | | |

### Figures

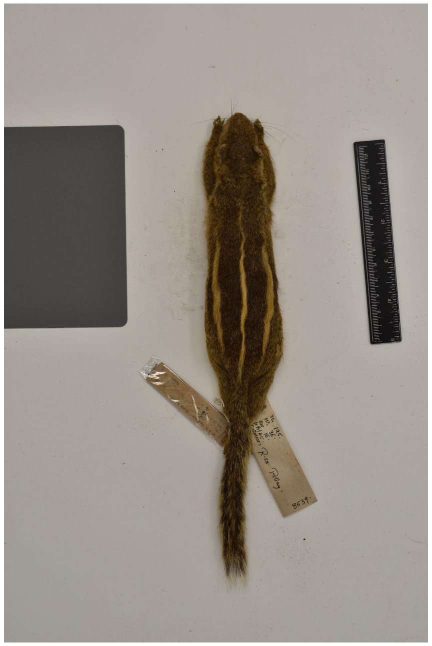

Figure S1. Image of a squirrel specimen with standards photographed under controlled light conditions

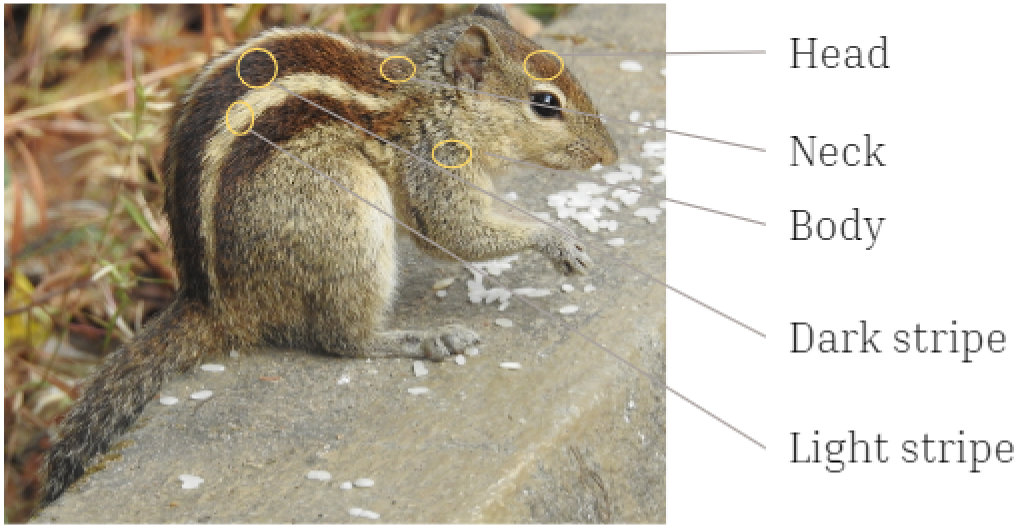

Figure S2. Positions of various body regions chosen for colour quantification

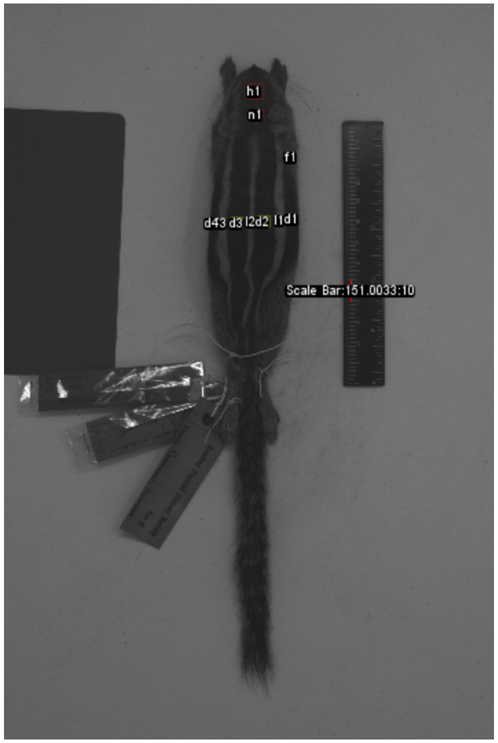

Figure S3. Multispectral image generated by micaToolbox in ImageJ

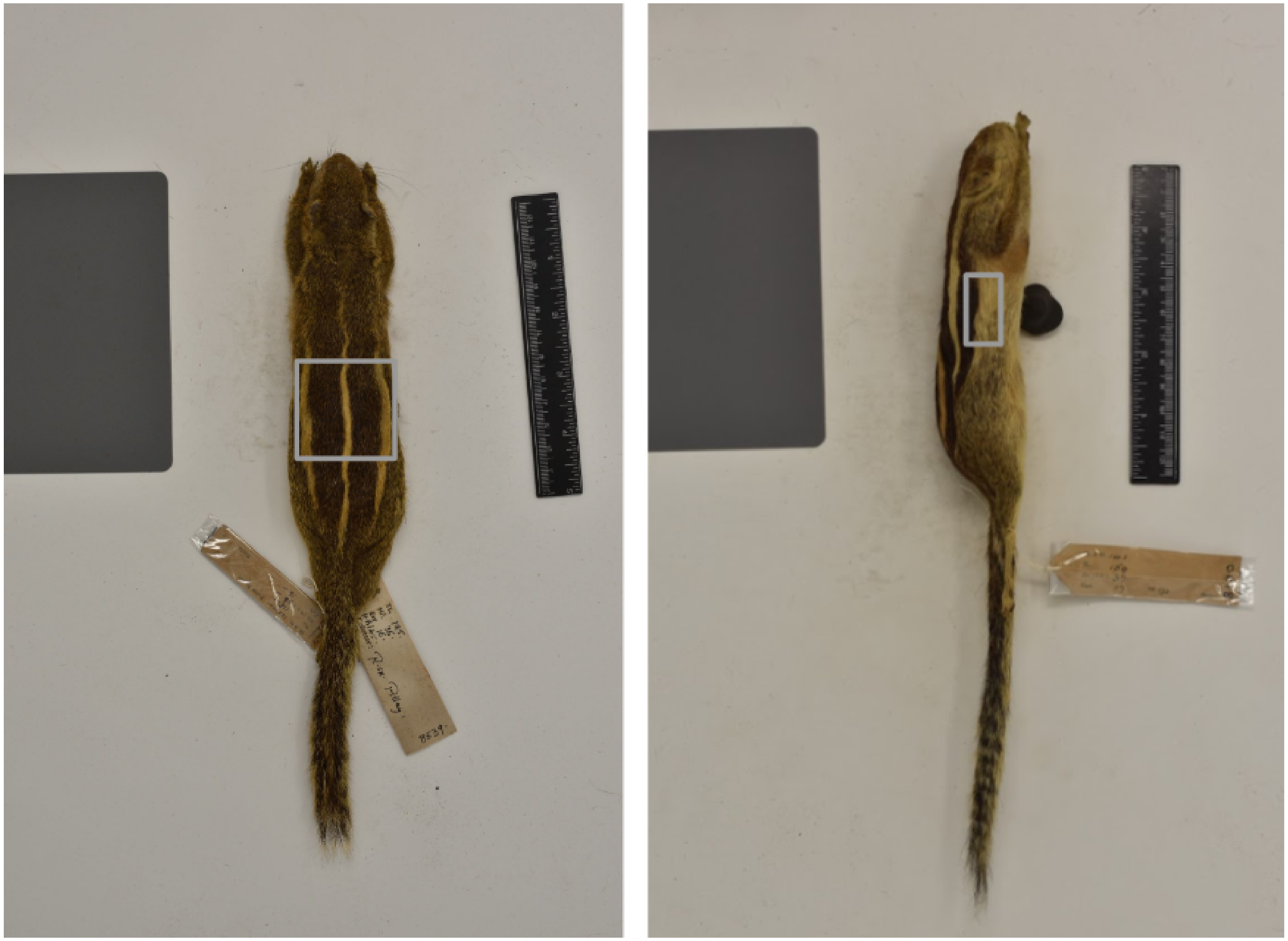

Figure S4. Selection of ROI for the dorsal (a) and lateral (b) stripe pattern analyses

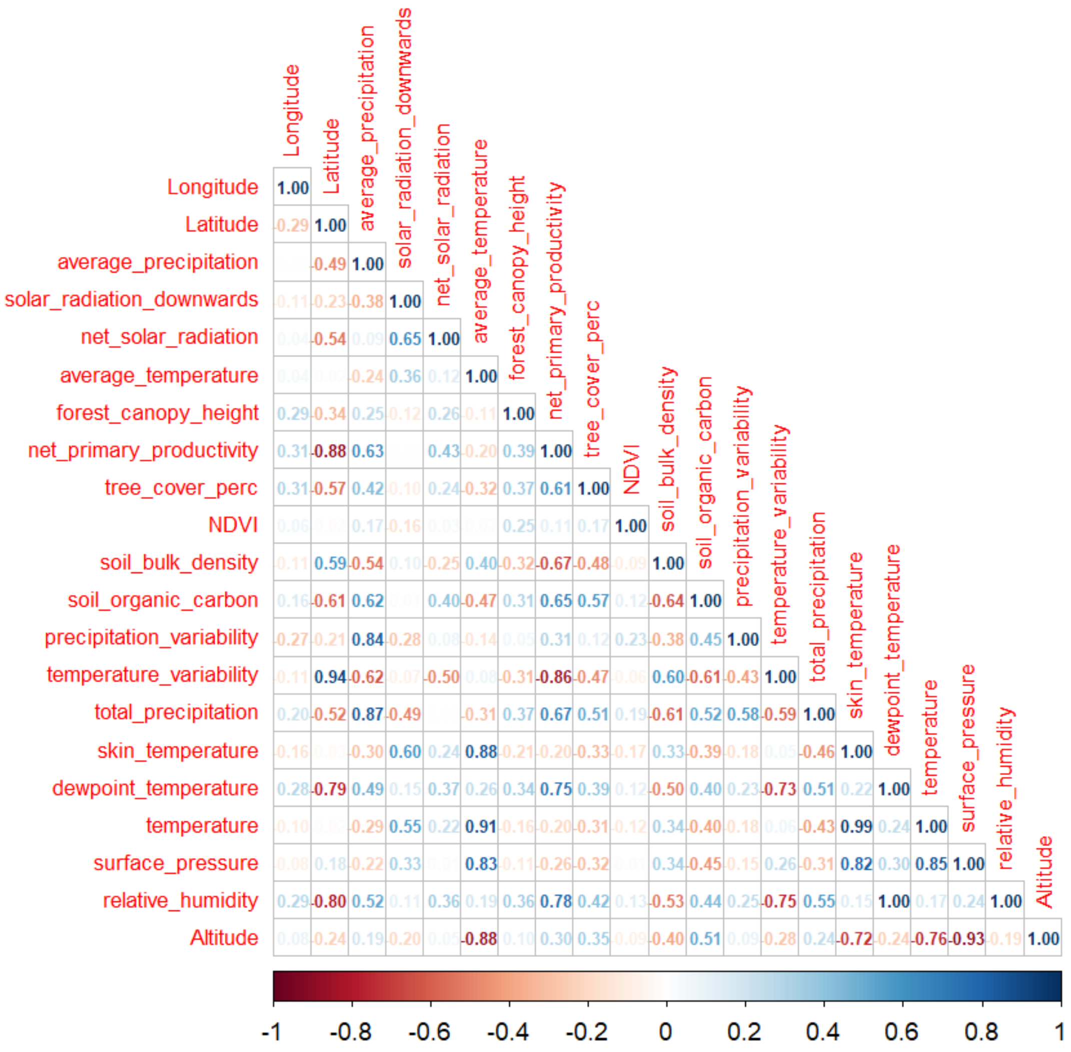

Figure S5. Correlation between environmental variables at squirrel locations

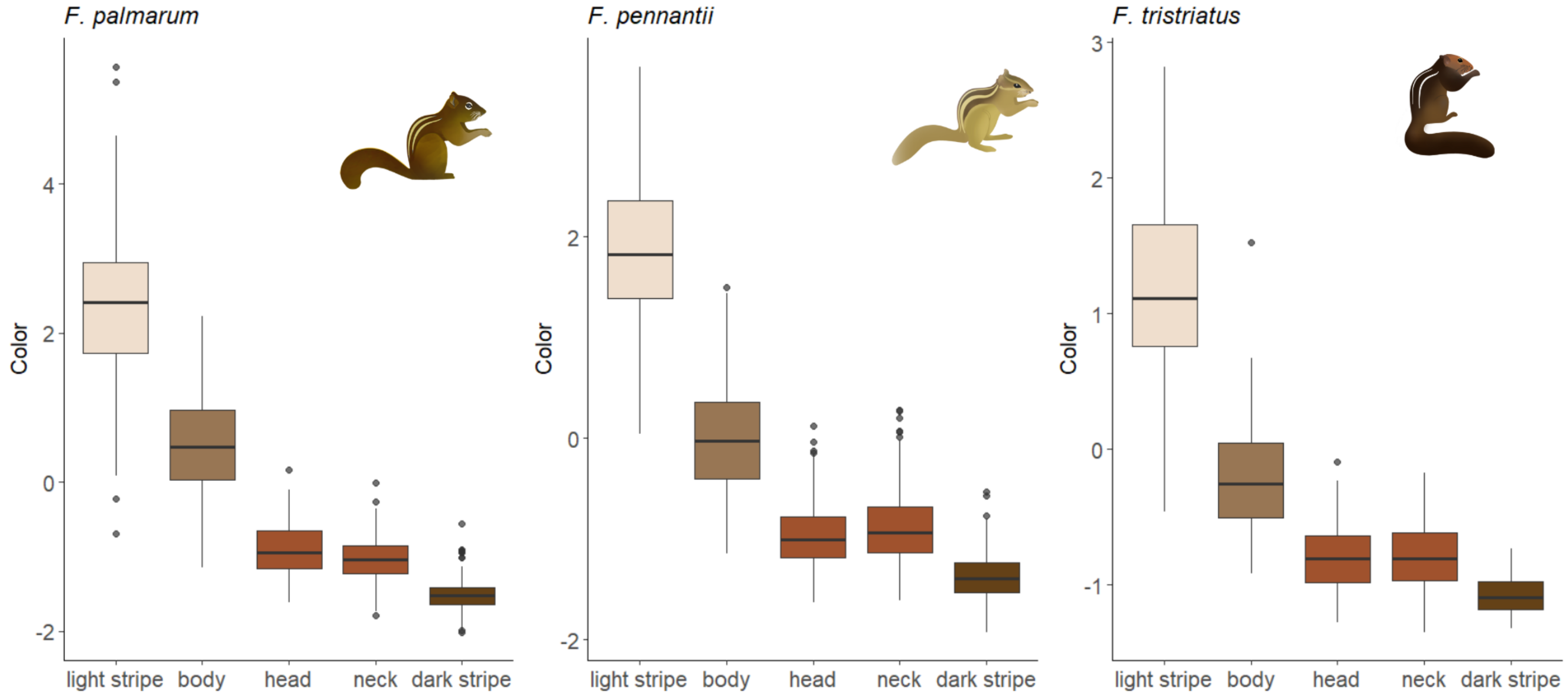

### Figure S6. Colour variation across body regions in each squirrel species.

Description: The colour values (y-axis) result from PC scores constructed from reflectance values of R, G, and B channels. Separate principal component analyses were performed for each species.

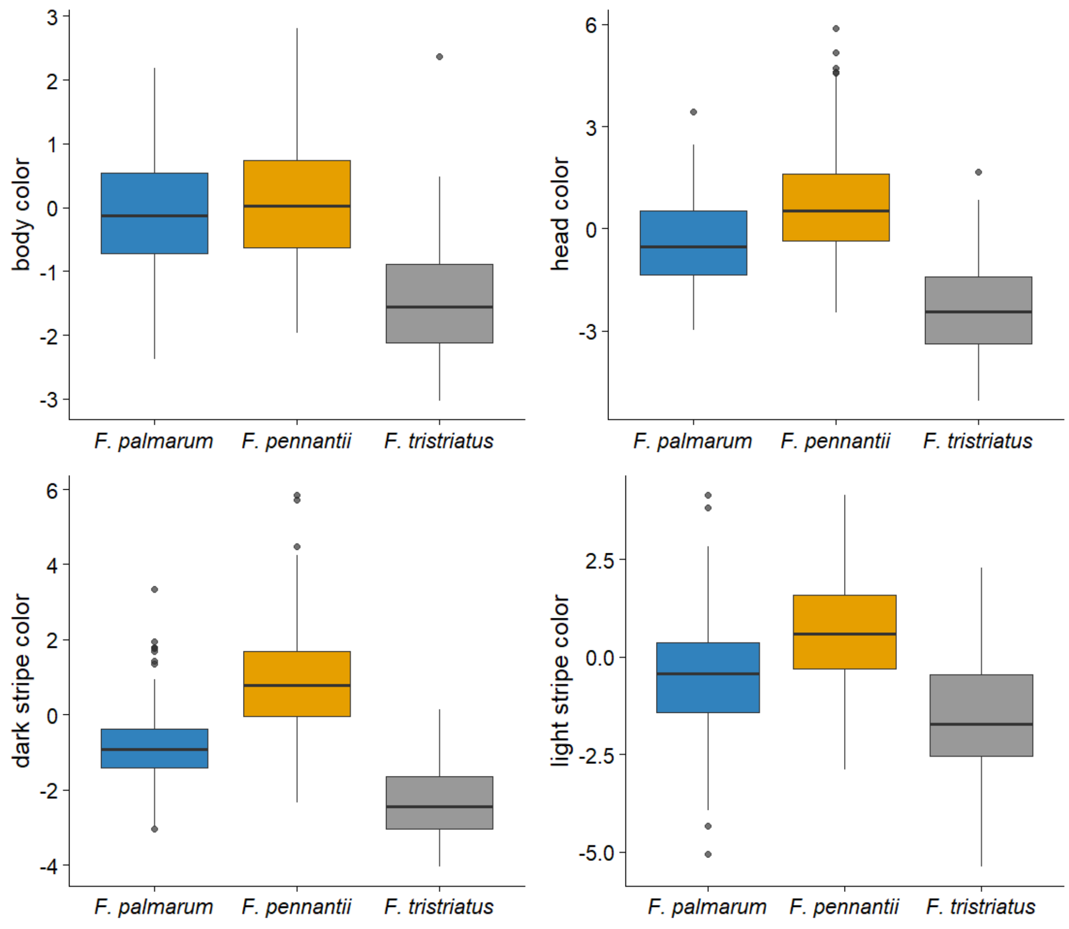

### Figure S7. Colour variation of a body region across species

Description: Colour (y-axis) values result from PC scores from reflectances in R, G, and B channels. Separate principal component analyses were performed for each body region across species.

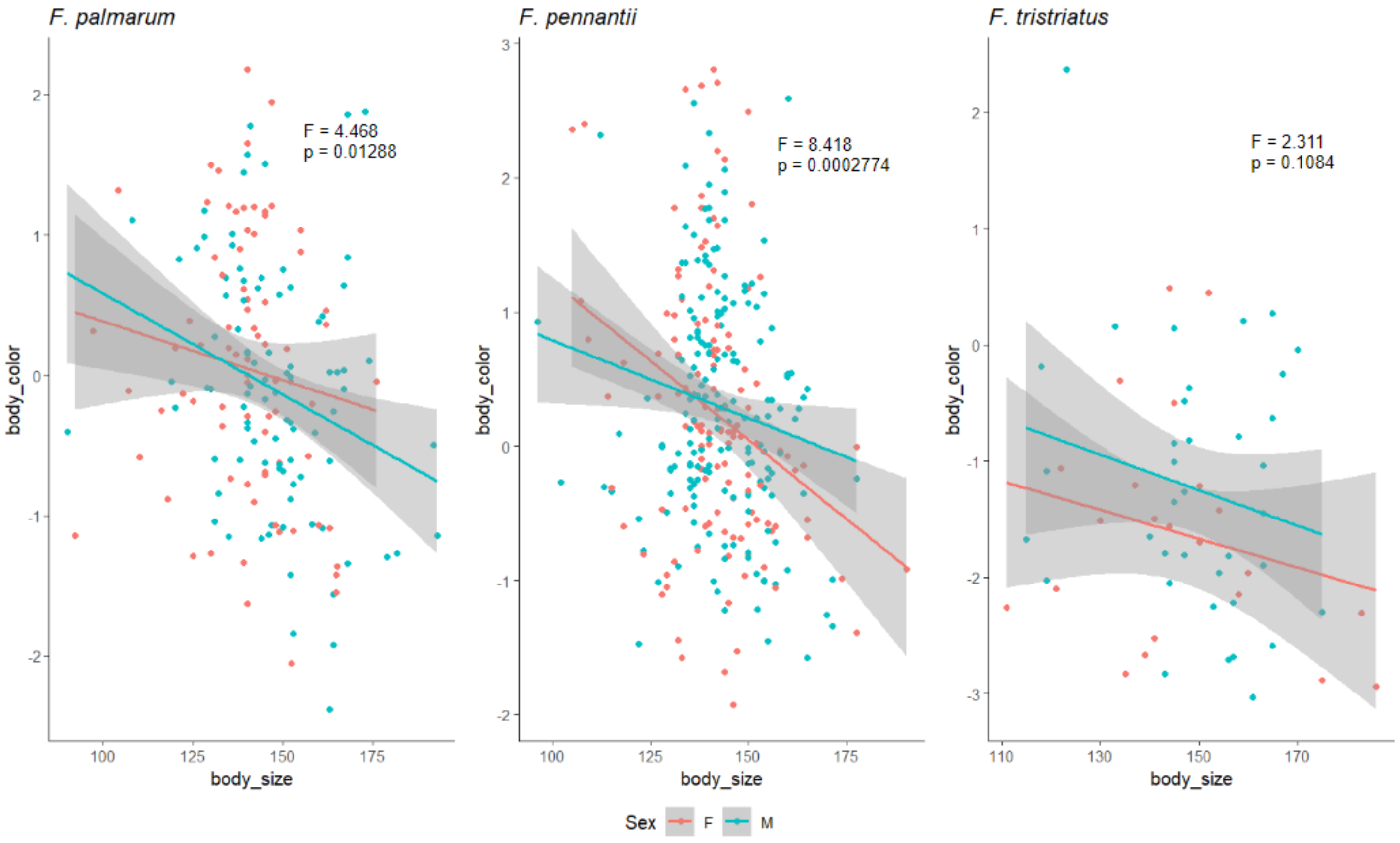

Figure S8. Relationship between coat colour and body size, grouped by sex

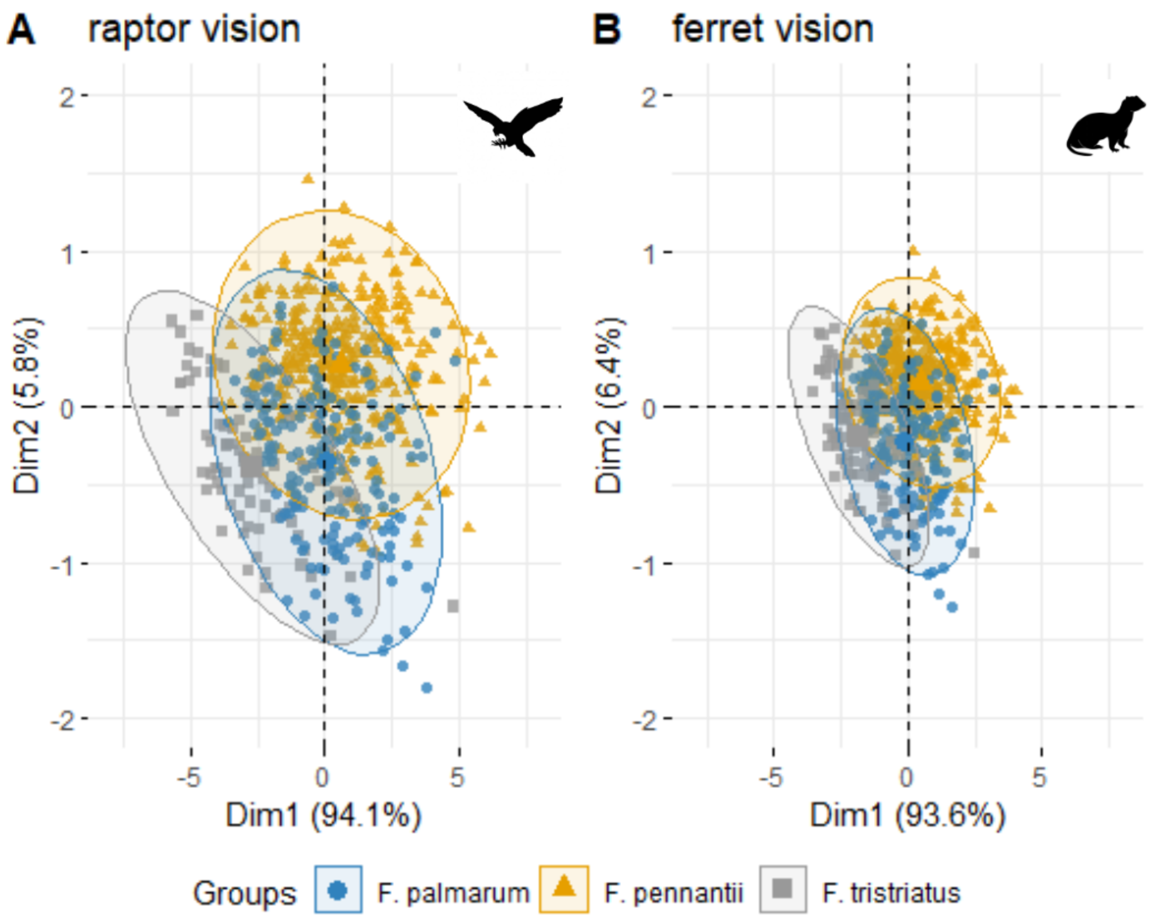

Figure S9. Variation of squirrel coat colour under ferret vs raptor visual systems

Description: PCA plot depicting the variation of coat colour among squirrel species as seen by the ferret and raptor predators.

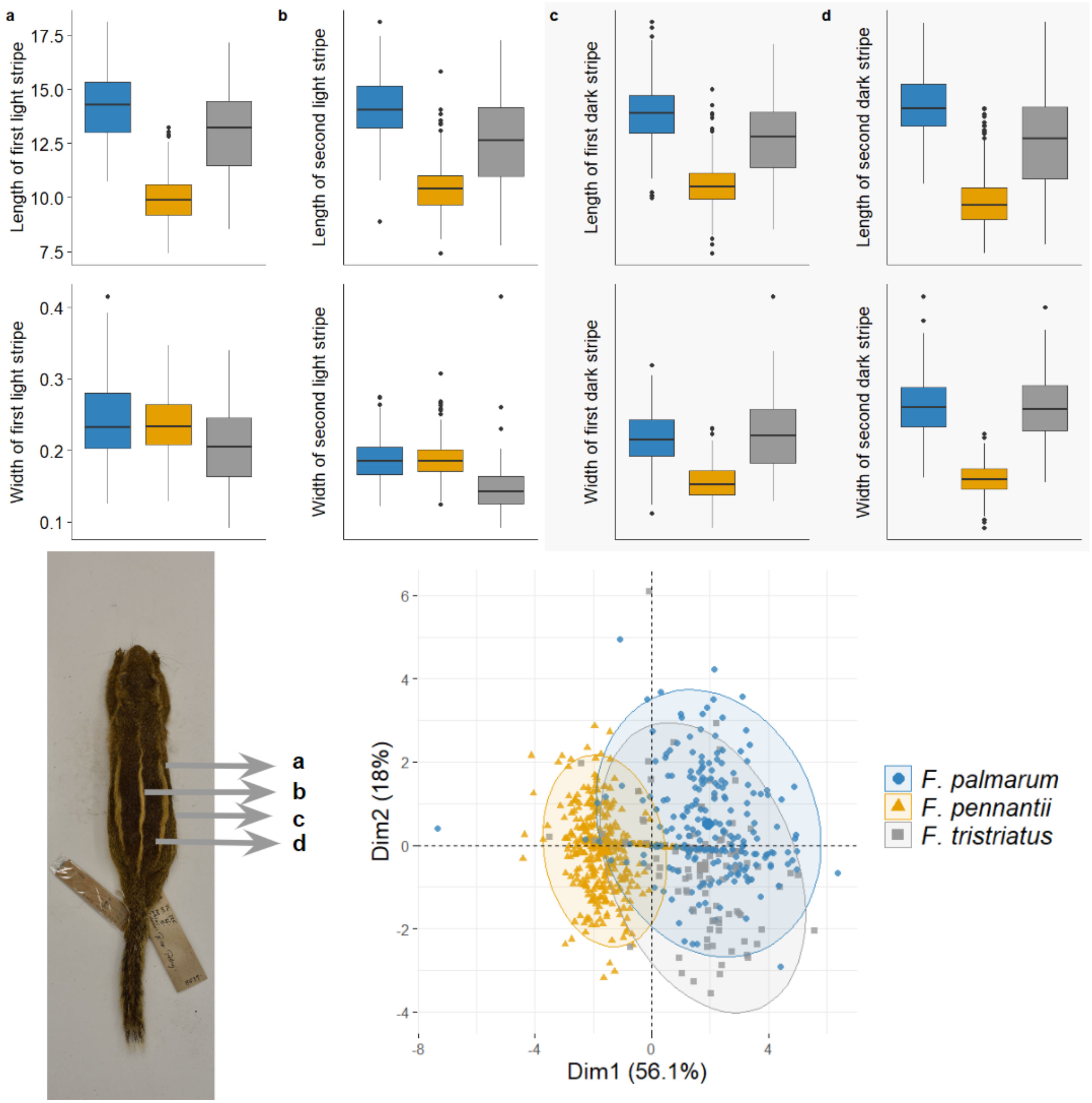
a
b
c
d
Length of first light stripe
Length of second light stripe
Length of first dark stripe
Length of second dark stripe
Width of first light stripe
Width of second light stripe
Width of first dark stripe
Width of second dark stripe
17.5
15.0
12.5
10.0
7.5
0.4
0.3
0.2
0.1
a
b
c
d
Dim2 (18%)
Dim1 (56.1%)
6
4
2
0
-2
-4
-8
-4
0
4
F. palmarum
F. pennantii
F. tristriatus

### Figure S10. Variation of stripe dimensions across species

Description: Interspecies variation of light and dark stripe dimensions

Raptor

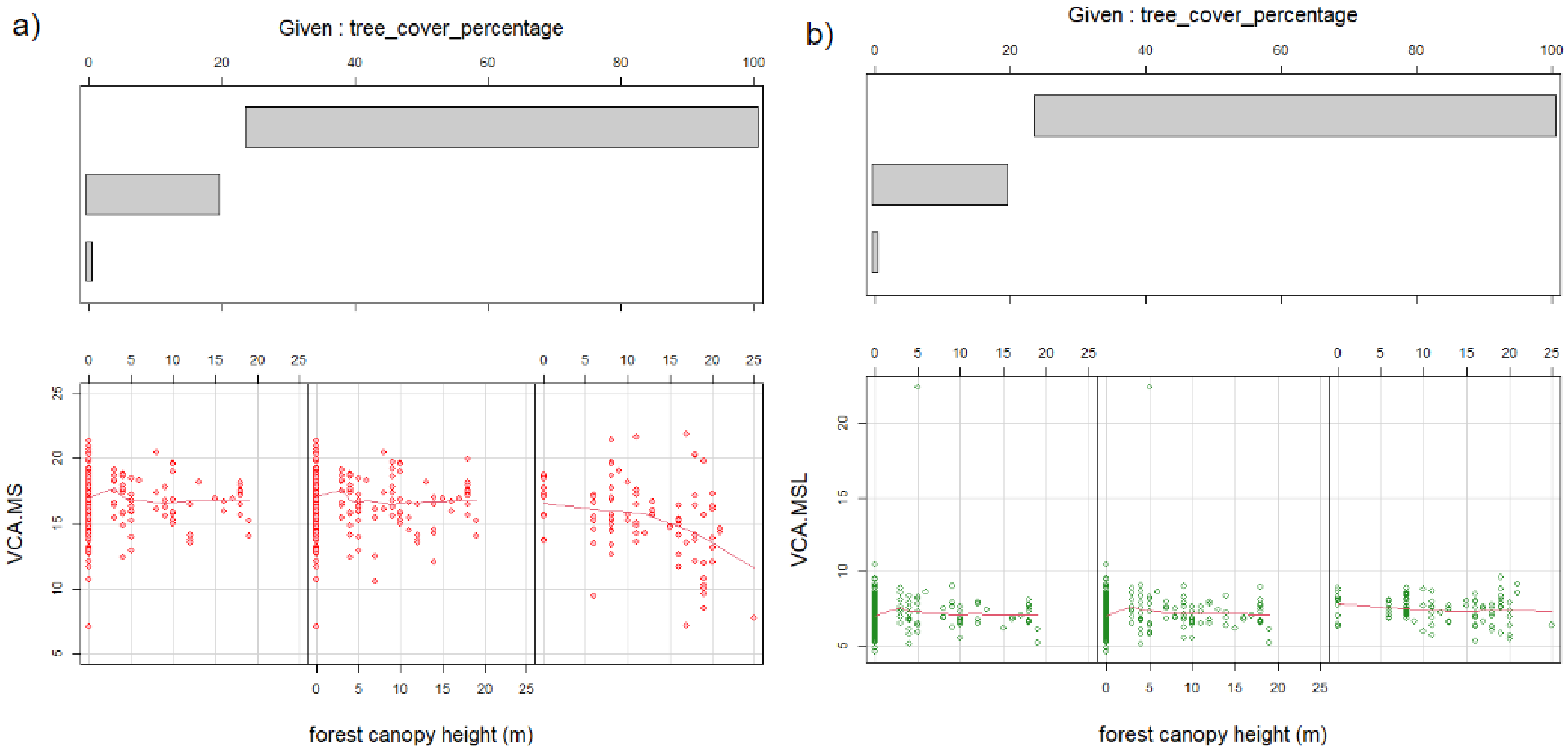

a)
Given : tree_cover_percentage
0
20
40
60
80
100
0
5
10
15
20
25
VCA.MS
forest canopy height (m)
b)
Given : tree_cover_percentage
VCA.MSL
forest canopy height (m)

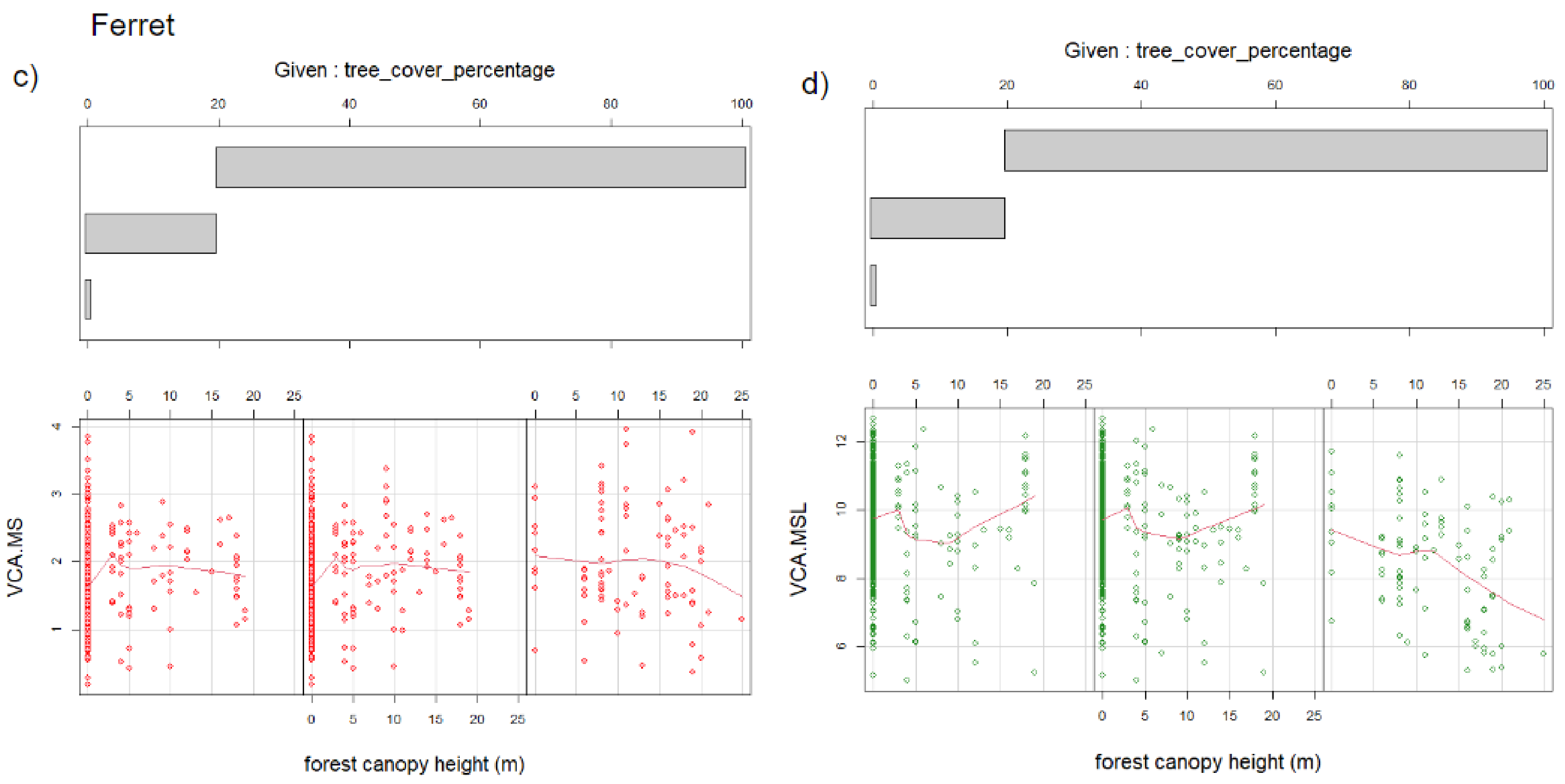

Figure S11. Variation of stripe contrasts with the interaction between tree cover and canopy height
